# Caging of membrane-to-cortex attachment proteins can trigger cellular symmetry breaking

**DOI:** 10.1101/2024.10.14.618153

**Authors:** Srishti Dar, Rubén Tesoro Moreno, Ivan Palaia, Anusha B. Gopalan, Zachary Gao Sun, Léanne Strauss, Richard R. Sprenger, Julio M. Belmonte, Sarah K. Foster, Michael Murrell, Christer S. Ejsing, Anđela Šarić, Maria Leptin, Alba Diz-Muñoz

## Abstract

To migrate, divide, and change shape, cells must regulate the mechanics of their periphery. The cell surface is a complex structure that consists of a thin, contractile cortical actin network tethered to the plasma membrane by specialized membrane-to-cortex attachment (MCA) proteins. This active and constantly fluctuating system maintains a delicate mechanochemical state which permits spontaneous polarization and shape change when needed. Combining *in silico*, *in vitro*, and *in vivo* experiments we show how membrane viscosity and MCA protein length regulate cortical dynamics. We reveal a novel mechanism whereby caging of linker proteins in the actin cortex allows for the amplification of small changes in these key parameters, leading to major alterations of cortical contractility. In cells, this mechanism alone gives rise to symmetry breaking phenomena, suggesting that local changes in lipid composition, in combination with the choice of MCA proteins, contribute to the regulation of cellular morphogenesis and function.

## INTRODUCTION

In biological systems, asymmetries exist at all levels – from molecules to large-scale polarities in tissues. At the cellular level, the establishment of shape polarities through symmetry breaking is fundamental to every cell type and complex biological process^1^. Cell division, growth and motion, and the establishment of complex cell shapes all require morphological asymmetries. A common way of breaking symmetry is through the presence of polarized external signals, with the most studied case of this being cell migration towards an attractant or away from a repellent^2,3^. However, a polarized signal is not always needed, for example in the case of T cells and neutrophils that can break symmetry in their shape in the presence of a homogeneous activator^4,5^. In such cases so-called “spontaneous” symmetry breaking occurs. On a theoretical level, small fluctuations or heterogeneities are sufficient to lead to symmetry breaking. In fact, a homogenous signal that strengthens interactions among components can reinforce feedback loops poising the system for symmetry breaking^6^.

The cell surface is the location of symmetry breaking for shape change phenomena. In animal cells, the cell surface is a composite interface comprising the plasma membrane, the actin cortex, and membrane-to-cortex linker attachment (MCA) proteins which connect them. In the context of cell shape changes, the most studied component of this system is the actin cortex, a dense network of dynamic and continuously turned-over actin filaments which lies beneath the membrane. This network is crosslinked by over a hundred different actin-binding proteins, including proteins which bundle and crosslink actin filaments, as well as contractile myosin-2 motors. Due to the strength of myosin-generated contractile forces^7^, the cortex has often been considered to be overwhelmingly ‘stronger’ than other components of the cell surface, with the membrane thought to participate in chemical signaling processes but passively following cortical deformations. Thus, in the context of symmetry breaking at the cell surface, local control of myosin contractility has dominated biophysical discussions^6,8^.

The biochemical feedback loops controlling actomyosin contractility, i.e. Rho/Rac/CDC42 signaling, undoubtedly play important roles in the generation and maintenance of cell polarity^9^. However, the membrane itself, despite being a fluid boundary, has physical properties which are highly regulated for their roles in cellular homeostasis. Membrane mechanics are usually discussed in terms of membrane tension and fluidity. Membrane tension arises from a combination of in-plane tension due to forces between lipids and friction due to the attachment to the underlying cortex^10,11^. On the other hand, membrane fluidity, which describes how freely components move about in the membrane, is primarily regulated via lipid composition. The plasma membrane is remarkably complex, containing over 300 different lipid species, as well as a host of peripheral and integral proteins and sugars^12,13^. Lipid composition is controlled both on a global level, for example to adapt to temperature fluctuations^14,15^, and on a local level for signaling^16,17^. Estimations based on membrane composition and fluorescent polarization signals suggest a 2-fold change in membrane fluidity between the apical and basolateral domains in epithelial cells^18–20^. Differences in membrane fluidity^21^ and lipid synthesis^22,23^ have also been observed between different stages of the cell cycle. In spite of the obvious importance of the proper regulation of membrane fluidity^24,25^, its study has been hampered by the lack of precise biophysical tools to modulate these properties in cells.

The third component of this composite, MCA proteins, connect the membrane to the actomyosin cortex. While in principle any protein that contains both filamentous-actin (F-actin) and lipid-binding domains can function as an MCA protein, the paradigmatic family are the ezrin-radixin-moesin (ERM) proteins^26,27^. ERMs have a conserved architecture (plasma membrane-binding and F-actin-binding domains connected via a flexible *α*-helical hinge) and biochemical regulation (phosphorylation)^28–30^. Phosphorylation in the F-actin-binding domain promotes a stable interaction between the plasma membrane and cortical F-actin^31^. ERM proteins play a crucial role in cellular symmetry breaking; phosphorylated ERMs are canonical polarity markers of the apical domain of epithelia (where ezrin was first identified^32^) and early mouse embryos^33^ and in the back of various migrating cells, including metastasizing cancer cells^34,35^ and T cells^36,37^. Consistent with the critical role of ERM proteins in immune cell motility, people with ezrin or moesin mutations present with immune deficiencies^38,39^. Various papers have hinted that in addition to a signaling role in recruiting the contractile components necessary to induce cortical deformations^40^ ERM proteins also play a structural role in symmetry breaking phenomena, for example by ‘coralling’ or inhibiting the motility of other proteins to enable the formation and maintenance of a polarized domain^27^. However, the underlying mechanism(s) remains to be elucidated.

Here, we examine symmetry breaking from the perspective of the cell surface as a composite interface. The synergy of biophysical characteristics of the different cell surface components leads to emergent properties which we will argue are employed by the cell to regulate complex phenomena during growth and function. We show that MCA proteins function as bidirectional mechanical integrators, transmitting mechanical information not only from the cortex to the membrane, but also from the membrane to the cortex. Thus, the biophysical properties of the membrane also regulate cortical mechanics via control of MCA protein mobility. Using a minimal actin cortex tethered to supported bilayers we show how variations in linker length and membrane fluidity alter cortical mechanics and contractility. This is due to length- and membrane viscosity-dependent protein ‘caging’ – whereby MCA proteins become trapped in the underlying actin cortex, which restricts their mobility and effectively stalls the cortical network. In cells, this gives rise to a new mechanism for symmetry breaking at the surface. Here, a positive feedback loop is established in which linker proteins caged in the cortical actin meshwork recruit more actin which in turn reinforces linker caging. Linker length and membrane fluidity are tunable amplifiers of this feedback loop. Such mechanisms highlight the importance of a materials science understanding of the cell surface composite. Furthermore, they do not depend on single genes but on the material properties that result from the combination of polymers and membranes, thus providing an inherent robustness to genetic instability. Overall, the biophysics of the cell surface composite reveals additional regulatory mechanisms which expand our understanding of the self-organizing capacity of active systems.

## RESULTS

### An *in vitro* reconstitution assay to test the role of a fluid boundary for actomyosin dynamics

Dissecting the crosstalk between the components of the cell surface composite is difficult in cells. Each of the components has myriad signaling roles and specific interventions to reliably manipulate cell surface mechanics are lacking. For example, membrane viscosity can be experimentally altered in cells by sequestering cholesterol using methyl-ß-cyclodextrin (MβCD). However, this has been shown to have additional effects such as shutting down numerous signaling pathways and changing actin levels^41–43^. We therefore began our experiments on an *in vitro* supported lipid bilayer system. We reasoned that alterations which affect cortical mechanics and contractility in such simple systems would be likely candidates for the formation of asymmetries in the intrinsically heterogeneous environment of the cell surface.

We first tested whether the biophysical properties of the membrane could influence cortical mechanics. To this end, we deposited membranes on silanized glass coverslips covalently attached to polyethylenglycol (PEG), which allows for full mobility of the lipids within the bilayer (**Figure 1A, first panel, Video S1**). Our starting fluid membrane condition comprised DOPC:DOPS:DGS-NTA(Ni^2+^):TxR at a ratio of 82.5:12:5:0.5 mol% (henceforth referred to as DOPC). We then used two different methods which have been previously shown to increase the membrane viscosity^44–48^, either by the addition of 30 mol% cholesterol (DOPC:DOPS:DGS-NTA(Ni^2+^):Chol:TxR at a ratio of 52.5:12:5:30:0.5 mol%; henceforth referred to as +Chol) or 60 mol% DOPE (DOPC:DOPS:DGS-NTA(Ni^2+^):DOPE:TxR at a ratio of 22.5:12:5:60:0.5 mol%; henceforth referred to as +DOPE). To validate the effect of these changes we imaged the membrane by confocal microscopy (**Figure 1B, first panel**) and quantified their fluorescence recovery after photobleaching (FRAP on Texas Red (TxR)) (**Figure 1C**). In agreement with previously published data^49–51^, we found that the addition of either 30% Chol or 60% DOPE leads to a 2.1-fold reduction in the diffusion coefficient of lipids without affecting the mobile fraction (% recovery) of lipids and caused no observable phase separation (**Figure 1D, E, Figure S1A, B**).

**Figure 1.**
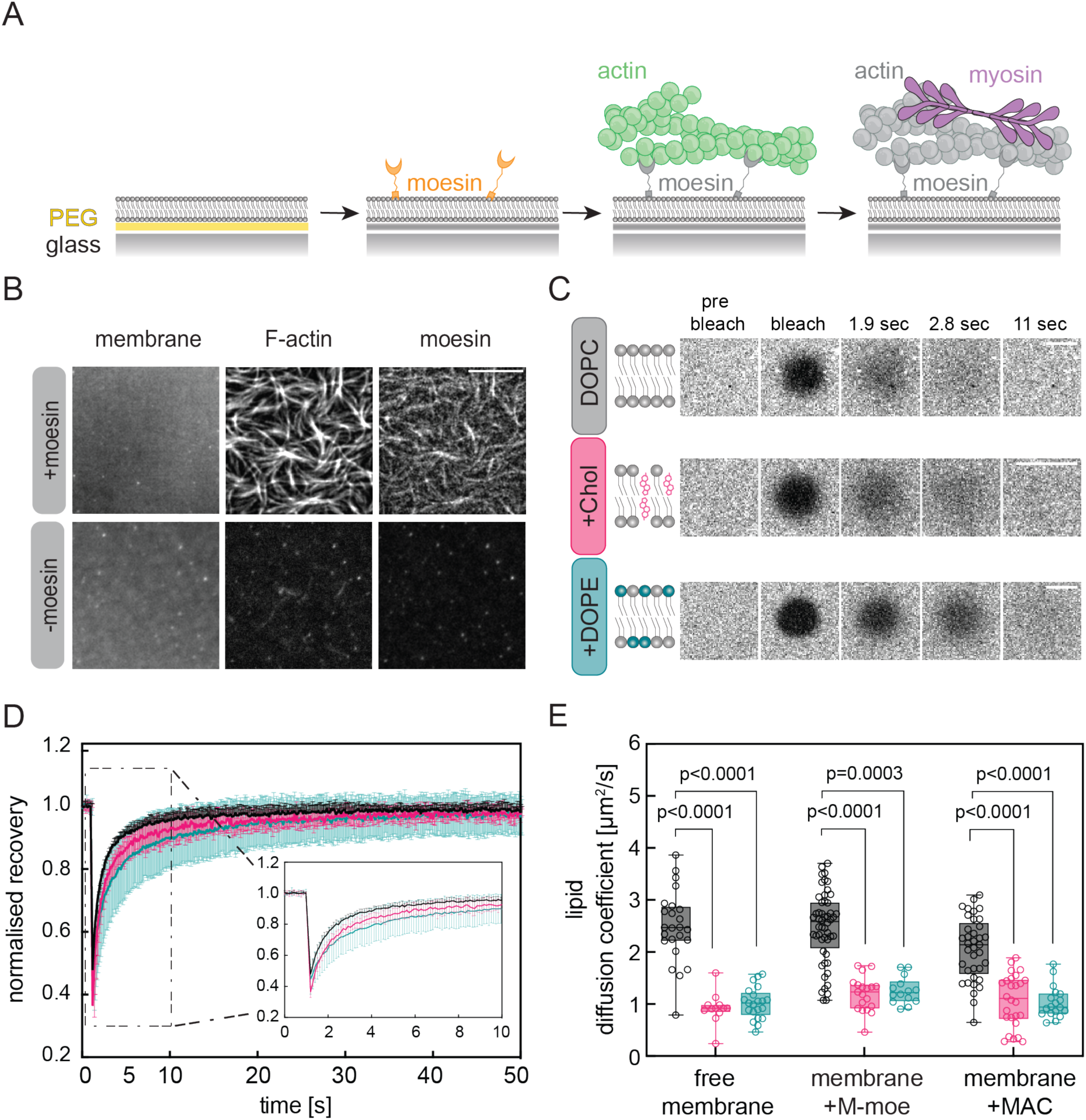
An *in vitro* reconstitution assay to address the role of coupling to a fluid boundary for actomyosin dynamics. A) Schematic of step-wise assembly of a reconstituted minimal actin cortex (MAC) on defined membranes. Molecules not drawn to scale. B) Representative epifluorescence images of reconstituted MACs with and without moesin. Membrane is labelled by Texas Red-labelled lipids, F-actin by Alexa 647-Phalloidin and moesin (M-moe) is GFP-tagged. Scale bar 10 µm. C) Schematic and representative confocal images of membranes containing 0.5 mol% Texas Red-labelled lipids. (n>5 curves/experiment N=3-5 experiments). Scale bars = 5 µm. D) Averaged traces from curves showing fluorescence recovery after photobleaching on DOPC (grey), +Chol (pink) or +DOPE (teal) free membranes; error bars indicate SD and bold line is the mean. E) Lipid diffusion coefficient of DOPC (grey), +Chol (pink) and +DOPE (teal) membranes. Solid line in box plots represents median. For significance: One-way ANOVA (Kruskal-Wallis multiple comparison test).

Having established a membrane template with tunable fluidity, we built-up complexity by adding a minimal actin cortex (MAC) tethered to the membrane by MCA proteins^52–54^ (**Figure 1A**). We first introduced a 6XHis-tagged moesin construct, which comprised the C-terminal actin binding domain (C-ERMAD) and its flexible alpha helix (275 amino acids; henceforth referred to as M-moe). The His-tag in this construct non-covalently binds the nickel-conjugated lipid in the bilayer and mediates coupling between the actin filaments and the membrane (**Figure 1B**). After washing the unbound protein, we flowed in a 2 µM solution of pre-polymerized actin filaments stabilized with phalloidin (0.5 µM), and incubated for 10 minutes before washing. This led to the formation of a dense network (i.e. start and end points of filaments could not be identified). We used FRAP to measure changes in membrane viscosity, by monitoring lipid diffusion coefficients (**Figure 1C-E)**. To confirm that the addition of a MAC does not create diffusion barriers by affecting the integrity of the PEG-cushioned bilayer we quantified the mobile fraction (% recovery) of lipids^26^ (**Figure S1A**) ^55,56^. We found that neither the addition of M-moe alone nor of a MAC to the supported lipid bilayer affected the increases in viscosity observed upon addition of cholesterol or DOPE.

It has been shown that changes in the cholesterol composition of the plasma membrane can lead to alteration in surface charge density, lipid packing, and membrane electrostatic potential, which could collectively enhance protein association with negatively charged membranes^57^. Different membrane conditions could thus lead to different binding of actin and/or linker protein. To ensure that protein content was consistent across our conditions, we added different concentrations of M-moe tagged N-terminally with GFP and quantified its fluorescence. In an independent experiment, we measured the fluorescence from Alexa 488-Phalloidin-labelled actin reconstituted on unlabelled M-moe-bound membranes. We selected final conditions such that once reconstituted all membranes displayed equal number of linker proteins and actin filaments before myosin addition (**Figure S1D-G**).

### Membrane viscosity can limit the ability of an actin network to be remodelled by myosin motors

Upon examination of the actin architecture in the *in vitro* system, we observed that the level of filament alignment varied according to the viscosity of the membrane, irrespective of incubation time (**Figure S2A**). Specifically, both the nematic ordering (**Figure 2A-C**) and apparent bundling (**Figure 2D**) decreased with increasing viscosity. Thus, membrane viscosity influences cortical architecture, even in the absence of cortical contractility.

**Figure 2.**
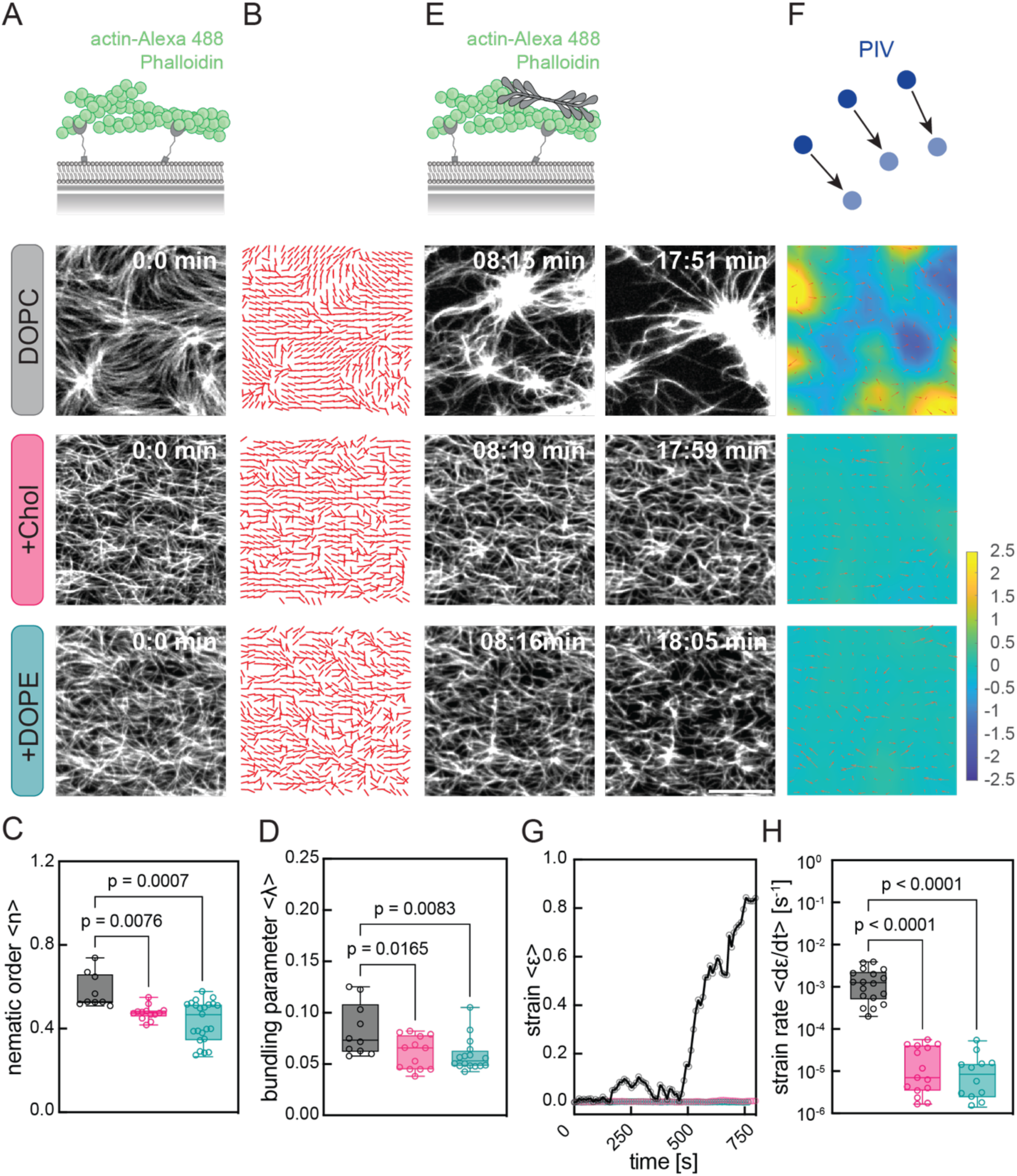
Membrane viscosity can limit the ability of an actin network to be remodelled by myosin motors. A) Schematic and representative images of Alexa 488-Phalloidin-labelled actin on DOPC, +Chol and +DOPE membranes. B) Skeletonization of representative nematic order on DOPC, +Chol and +DOPE membranes. Quantification of the nematic order (C) and apparent bundling (D) on DOPC (grey), +Chol (pink) and +DOPE (teal) membranes (n=3-7 images/experiment; N=3 experiments). E) Schematic with myosin added and representative time-series of actin remodelling on DOPC, +Chol and +DOPE membranes. F) Schematic of PIV analysis and representative vector fields. Quantification of mean strain over time (G) and strain rate (H) on DOPC (grey), +Chol (pink) and +DOPE (teal) membranes (n=2-4 ROIs/experiment; from at least N=3 experiments). Scale bar in E = 10 µm. Time stamp represents the time from myosin addition in min:sec. Calibration bar shows strain distribution. Solid line in box plots represents median. For significance: One-way ANOVA (Kruskal-Wallis multiple comparison test). See also Figures S1 and S2, and Videos S2, S3 and S4.

Next, we assessed the remodelling activity of myosin on the MACs. Upon addition of pre-polymerized muscle myosin minifilaments (100 nM) (**Figure 2E** and **S2B**) myosin is able to remodel the actin network into asters only on DOPC membranes. On both +Chol and +DOPE membranes the addition of myosin had little or no effect on the structure of the actin cortex (**Figure 2E-H**, **Videos S2-4**). We quantified the deformation of the actin network in response to myosin addition by Particle Image Velocimetry (PIV)^58^. This allowed us to determine the average actin displacement (mean strain, ε) over time, and the strain rate, dε/dt (**Figure 2F-H**, **Methods)**. In more fluid (DOPC) membranes, we observed the establishment of a consistent inward contractile flow within minutes following addition of myosin. Significant network deformations occur as the system restructures into asters, as has been previously reported^58–64^. This rapid reorganization and compression of F-actin lead to two orders of magnitude increase in strains and strain rates. Conversely, in membranes with cholesterol or DOPE, i.e. under high viscosity conditions, no change was observed (**Figure 2F-H**). This was despite these networks having comparable actin density (**Figure S1G**) and filament length distributions (**Figure S2C, D**). Together, these results show that a two-fold increase in membrane viscosity is sufficient to dramatically alter cortical dynamics and induce a switch to an arrested F-actin network that is able to resist the propagation of myosin-generated contractile forces. Thus, alterations in membrane viscosity provide an elegant and powerful way to alter cortical mechanics.

### Actin-bound MCA linker diffusion is sensitive to membrane viscosity

How can a two-fold change in the diffusion coefficient of the membrane translate to a 100-fold change in cortical strain rate? To understand how membrane viscosity regulates contractility, and rule out the contribution of different actin architectures (**Figure 2B-D**) to network arrest (**Figure 2F-H**), we turned to simulations (**Figure 3A, B**). We used a minimal model based on polymers composed of volume-excluded spherical beads, connected by springs. Actin was represented as long polar polymers. Myosin was modelled as a two-bead polymer, whose heads bind and unbind from actin filaments and directionally walk along them, dwelling on their barbed ends^65,66^. Based on previous simulations of actin filaments on negatively charged membranes^73^, similar to our *in vitro* conditions, we introduced a gap between the plane (membrane) and the cortex. Such a gap also exists in cells^74,75^, where additional factors such as protein crowding and binding kinetics of various linker proteins to a dynamic cortex are also at play. Linkers were four-bead polymers tethered to a fluid horizontal plane from one end, binding and unbinding from actin filaments in the other end and diffusing in the x-y plane (see **Methods** for details). We assessed the behaviour of these networks before (**Figure 3A**) and after the addition of myosin (**Figure 3B**), for different viscosities of the medium, trying to recapitulate the impact of the two-fold membrane viscosity increase observed in experiments (**Figure 1E)** but with identical initial actin architecture.

**Figure 3.**
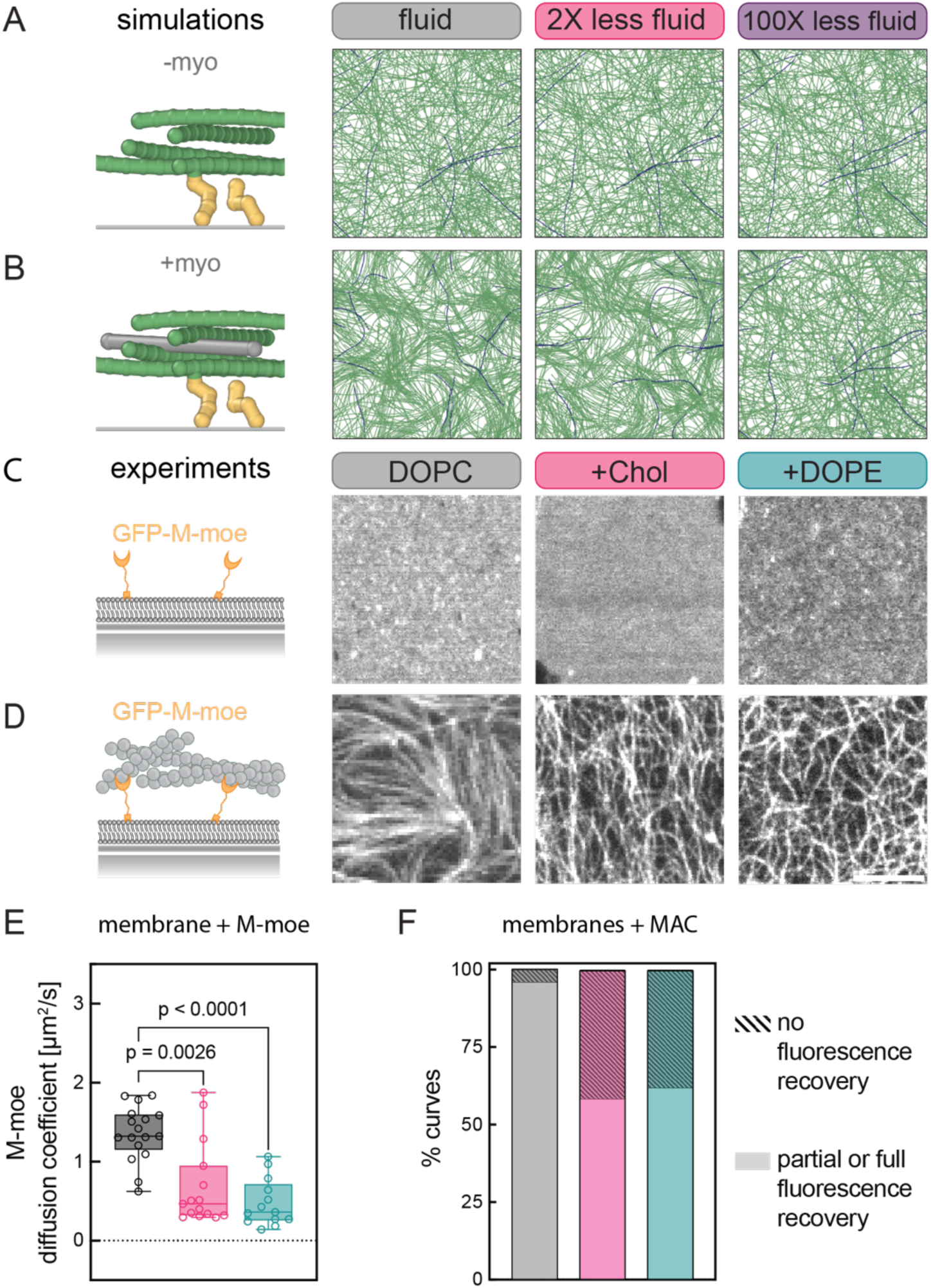
MCA linker diffusion does not scale with membrane viscosity when bound to an actin network. A-B) Left: Schematic of actin filaments, linkers, and myosin (lateral view). Right: Snapshots of simulations before (A) and after (B) myosin addition (top view). Some actin filaments are coloured in blue to ease interpretation. Medium viscosity decreases from left to right, as indicated. C-D) Representative epifluorescence images of GFP-M-moe recruited on membranes without (C) and with (D) F-actin (unlabelled). E) FRAP-derived diffusion coefficients of GFP-M-moe on DOPC (grey), +Chol (pink) and +DOPE (teal) membranes before actin addition (n≥5 curves/experiment, N=4,4,3 experiments). Solid line in box plot represents median and error bars indicate SD. F) Curves from FRAP experiments on GFP-M-moe after actin addition on DOPC (grey), +Chol (pink) and +DOPE (teal) are classified as to whether or not fluorescence recovered. If fluorescence partially recovered, the curve is classified as recovered. Scale bar in D = 5 µm. Solid line in box plots represents median. For significance: One-way ANOVA (Kruskal-Wallis multiple comparison test). See also Figure S3.

A two-fold reduction in medium viscosity, comparable to the one observed in our *in vitro* system, did not alter contraction in simulations (**Figure 3B**, left *vs.* centre, **3G**, **Videos S5 center**). A two orders of magnitude increase in viscosity recreated an arrest in contractility (**Figure 3B**, left *vs*. right, **Video S5 right**); however, this is inconsistent with our findings and would require non-physiological membrane viscosities.

Thus, a two-fold change in viscosity is insufficient to cause arrested networks, and some other mechanism must be at play. In our simple, signalling-independent *in vitro* system, such an effect must arise from the combination of mechanical properties of the different components. We reasoned that the information from the membrane must be transmitted to the cortex mechanically, via the MCA linker proteins. We therefore examined the mobility of the MCA proteins by FRAP, both before and after the addition of actin (**Figure 3C-F** and **S3**).

When bound only to lipids, M-moe showed a largely uniform distribution on membranes (**Figure 3C**), with a lateral mobility approximately 2- and 2.7-fold lower on +Chol and +DOPE membranes than on DOPC alone (**Figure 3E**). Fluorescence recovery was observed for all curves (**Figure S3-D**). However, the addition of actin led to a dramatic change in linker distribution from homogeneous to filamentous, mirroring F-actin (**Figure 1B, 3D**)^67^. Additionally, the diffusion of the linker was significantly affected by membrane viscosity when bound to an F-actin network. In the low viscosity DOPC condition, over 96% of FRAP curves showed complete fluorescence recovery (**Figure 3F**); however, many curves in the high viscosity conditions failed to recover (41.7% on +Chol and 38.1% on +DOPE compared to 4% on DOPC; **Figure 3F** and **S3-E**). The high fluorescence recovery on DOPC membranes indicates that the failure to recover is not simply due to the presence of bound F-actin, and suggests that some other mechanism must contribute to preventing the diffusion of a large population of linkers in the +Chol and +DOPE conditions.

### Membrane viscosity and the size of MCA proteins are rheostats that regulate cortical dynamics

Complete stalling of myosin-mediated contractility and disruption of linker motility were unexpected outcomes, both when compared to the magnitude of the causative disruption (a two-fold change in membrane viscosity), and in terms of their potential impact on cell surface mechanics in cells. To better understand this phenomenon, we sought ways to rescue myosin contractility in the higher viscosity condition. Briefly, flowing in excess myosin (200 nM) on the fully-connected arrested networks did not rescue their contractility (*data not shown*).

Other researchers have also observed contractility differences and/or stalling in similar minimal systems. Vogel *et al.* and Köster *et al.* observed contractility even with high membrane cholesterol content^68,69^. In contrast, Liebe *et al*. observed partial stalling of contractility even on perfectly fluid membranes^70^. What the former studies have in common is the use of very small artificial linker proteins. In part due to the difficultly of reconstituting and expressing full length ERM proteins in such a system, most studies to date have used similar solutions (single actin binding domains of fimbrin^59,71^, ezrin^69^, α-actinin^66^, or VCA^72^; or biotinylated lipids bound via neutravidin to biotinylated actin^64,68^). By contrast, Liebe *et al.,* used the full length ezrin protein in a fully connected network. We therefore wondered whether the length of the MCA linker was a critical factor, and whether we could tune network contractility by adjusting MCA protein length.

To test this, we generated two new constructs, one longer and one shorter than the one we had used so far. Specifically, for the short construct we used only the actin binding domain of moesin (C-ERMAD, 75 aa; henceforth referred to as S-moe) and for the long construct we used the full length moesin protein (577 aa; henceforth referred to as L-moe) (**Figure 4A, Figure S4A-C**). The influence of linker length on the actin architecture was readily apparent when we added F-actin to DOPC or +Chol membranes (**Figure 4B, left, C-D**). Systems with S-moe always had a high nematic order and bundling regardless of membrane viscosity, with the order and bundling parameters reaching similar values as those seen for M-moe on fluid DOPC (**Figure 4C-D**). This is consistent with previous studies using short linkers that do not report arrested networks^53,54^. For M-moe we found a dependence on viscosity: on fluid DOPC, the network had a high nematic order and bunding, resembling the S-moe networks, whereas on +Chol membranes it was more similar to L-moe.

**Figure 4.**
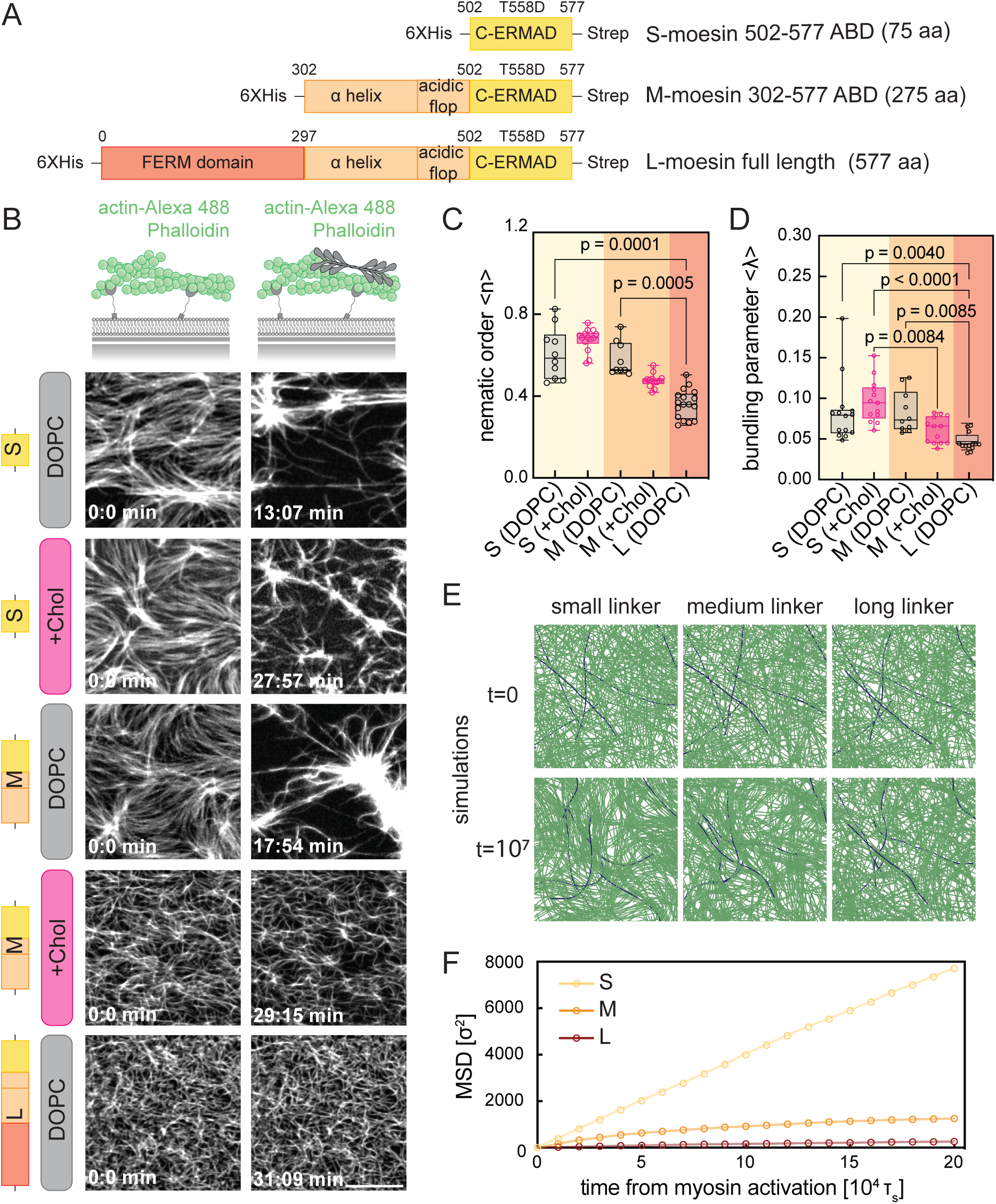
Membrane viscosity and the size of MCA proteins are additional rheostats that synergistically regulate cortical dynamics. A) Domain organization of moesin constructs. S, M and L represent the different truncated constructs of moesin. Numbers on top of domans represent amino acid count. B) Schematic and representative epifluorescence time-lapse images of fluorescently-labelled actin recruited on DOPC and +Chol membranes with S, M-, or L-moe constructs before (left) and after (right) myosin addition. Quantification of the nematic order (C) and bundling analysis (D) (n≥3 images/experiment; N=3). E) Snapshots from simulations of MACs with linkers of increasing length (S, M and L), before and after myosin addition. F) Mean square displacement (MSD) of actin filaments from simulations in (E). Scale bar in B= 10 µm. Time stamps min:sec. Solid line in box plot represents median. For significance: One-way ANOVA (Kruskal-Wallis multiple comparison test). See also Figure S4.

To observe the effect on contractility, we flowed in myosin and acquired movies until no further change in the actin architecture was observed. In the S-moe condition, myosin was able to remodel the cortex and induce aster formation even on the high viscosity +Chol membranes. However, remodelling was twice as fast on DOPC as on +Chol membranes (**Figure 4B, right**). In contrast, no remodelling occurred in the L-moe condition, even on the DOPC membranes (**Figure 4B, right**). To clarify the role of linker length *vs.* that of actin architecture, we once again turned to simulations of our MACs (**Figure 4E-F**). We started our simulations at equal actin concentration with randomly positioned filaments. The mobility of actin filaments upon myosin addition decreased with increasing linker size: the network was fluid for short (4 beads) and medium (9 beads) linkers and static for long ones (20 beads) (**Video S8,** see **Methods** for details). We quantified this by the actin mean square displacement, markedly suppressed for longer linkers (**Figure 4F**). The L-moe containing composites were thus completely arrested both *in vitro* and *in silico* (**Figure 4B, left** and **4E, bottom**), regardless of initial actin architecture. Thus, linker length influences the ability and timing of myosin-induced membrane remodelling, with medium-length linkers occupying a ‘sweet spot’ in which the capacity of myosin to remodel the membrane can be tuned by alterations in membrane viscosity.

### Protein caging determines MCA linker mobility and the ability of an actin network to be remodelled by myosin motors

To better understand the mechanism by which membrane viscosity and linker length might modulate actin architecture and contractility, we developed a new minimal computational model. In this model we removed myosin and froze actin filaments, allowing us to disentangle linker dynamics from cortex contractility. As in the previous model, linkers were tethered on one end to a horizontal plane representing the membrane, while the other end was free to bind and unbind actin. The cortex was simulated as rigid, immobile filaments. We again imposed a gap between the cortex and the plane (membrane). In this model, the membrane and the gap were coarse-grained as a region of lower viscosity than that of the cortex (**Figure 5A**, we label the 6-bead linker as S, 9-bead as M, and 24-bead as L, see **Methods** for details).

**Figure 5.**
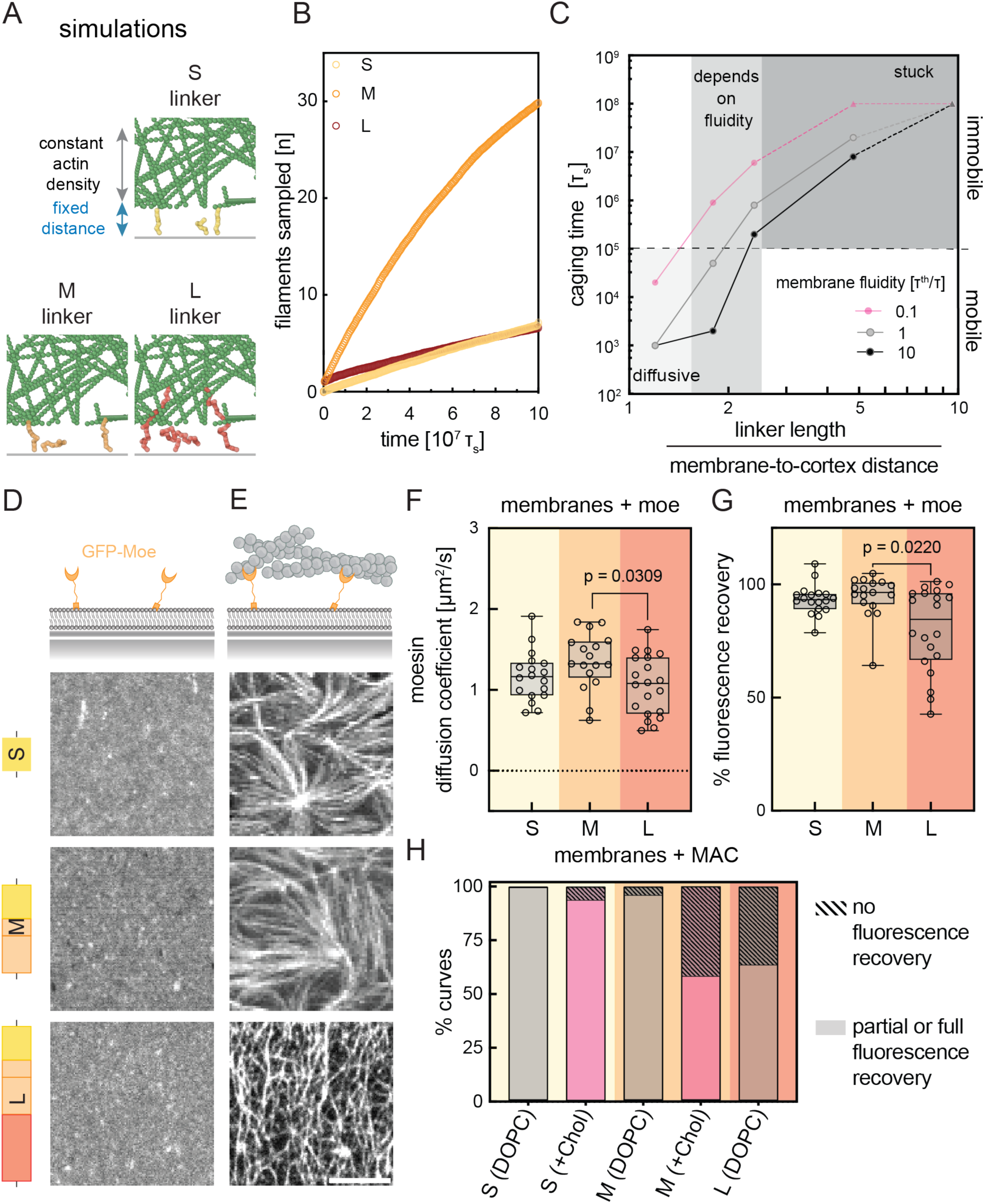
MCA protein caging determines its mobility and the ability of an actin network to be remodelled by myosin motors. A) Snapshots from simulations of linkers of various lengths (S, M and L), binding and unbinding to static filaments in the cortex. B) Cumulative number of actin filaments sampled over time by linkers of different length. C) Mean caging time as a function of linker length under different fluidity conditions. The dashed horizontal line represents a hypothetical observation time at which short linkers would appear mobile and long ones immobile irrespective of membrane fluidity, while intermediate linker lengths would only appear mobile on more fluid membranes. This is the situation depicted in Figure 4B. See **Methods** for details. D-E) Representative schematics and images of GFP- labelled S-, M-, and L-moe constructs recruited on membranes before (D) and after F-actin (E) addition. F) Diffusion coefficients and % fluorescence recovery (G) of GFP-labelled S-, M-, and L-moe constructs before F-actin addition on DOPC membranes (n≥5 curves/experiment, N=3 experiments). H) Curves from FRAP experiments with GFP-labelled S-, M-, and L-moe on DOPC (grey) and +Chol (pink) membranes are classified as to whether or not fluorescence recovered. If fluorescence partially recovered, the curve is classified as recovered (4 <N>10 curves/experiment, N=3,4,3 experiments). Scale bar in D = 5 µm. Solid line in box plots represents median. For significance: One-way ANOVA (Kruskal-Wallis multiple comparison test). See also Figure S5 and Video S7.

We quantified the frequency with which a linker changes binding partners (**Figure 5B)** and used the linker mean square displacement to compute the “caging time”, defined as the average time a linker is confined in a region before its motion becomes diffusive again (**Figure 5C, S8F**). We adopt the idea of caging from glass and polymer physics^76–78^ ^79^ as has been done before for biological systems^58^. Our simulations showed that there is a maximum exploration of the cortical interface at medium linker lengths (**Figure 5B).** Short linkers only bound few filaments close to the membrane due to the entropic penalty which was incurred upon full extension, and long ones became entangled within the cortical network leading to frequent rebinding to the same filament or to filaments within the same cortex mesh. The caging time increased with linker length, spanning 5 orders of magnitude for only a 4-fold change in linker length (**Figure 5C**). As expected, increasing binding energy (**Figure S8E**) or actin density in the cortex (**Figure S8D**) boosted this effect. This confirmed our prediction that caging resulted from the combination of actin-linker binding and steric entanglement, which together restricts the movement of the linkers. Furthermore, an increase in membrane viscosity (reducing linker mobility) increased caging times (**Figure 5C**), in agreement with our observation that viscosity changes cause networks containing M-moe to transition between contractile and arrested states (**Figure 2E-H** and **4B**).

To test the predictions from our simulations, we used FRAP of fluorescently labelled linkers to quantify the mobility of the S-, M- and L-moe on DOPC before and after actin addition (**Figure 5D, E**). All constructs had a similar diffusion coefficient on DOPC membranes before engagement with F-actin (**Figure 5F, G**). Once actin was added, fluorescence recovered only for the S-moe and M-moe constructs on DOPC and for the S-moe construct on cholesterol membranes, fully matching the conditions where contractility was observed (**Figure 4B, 5H and S5**). For M-moe on +Chol and L-moe on DOPC we found that 41.6 % and 36.4% of the curves failed to recover. We observed a positive relation between lateral mobility of linkers as a function of their length and percentage recovery upon actin engagement (**Figure S5F**), suggesting a membrane viscosity-independent role for linker length in regulating cortical mechanics.

### MCA linker protein caging leads to symmetry breaking in cells

Our *in vitro* system is highly simplified and homogeneous when compared to the complexity of the cell surface: lipids and proteins are homogeneously distributed, for example, and no actin remodelers or crosslinkers (aside from myosin) are present. Heterogeneities in linker distribution and membrane mechanics in cells could thus translate to uneven patterns of cortical mechanics. We therefore wondered in what way the *in vivo* implications of linker caging would differ from the homogeneous alterations in contractility patterns that we observed *in vitro*.

To explore this, we first overexpressed phosphomimetic moesin (CA-Moe; T558D) labelled with mCherry under a doxycycline inducible promoter in NIH3T3 fibroblast cells. We plated cells on round fibronectin micropatterns to restrict cell spreading and induce the formation of a uniform cell cortex (**Figure 6A**). This expression alone was sufficient to induce symmetry breaking in both the actomyosin cortex and linker distribution. Specifically, both actin and linker accumulated in a small patch of the surface; in extreme cases this led to a change in cell curvature (**Figure 6B-D**).

**Figure 6.**
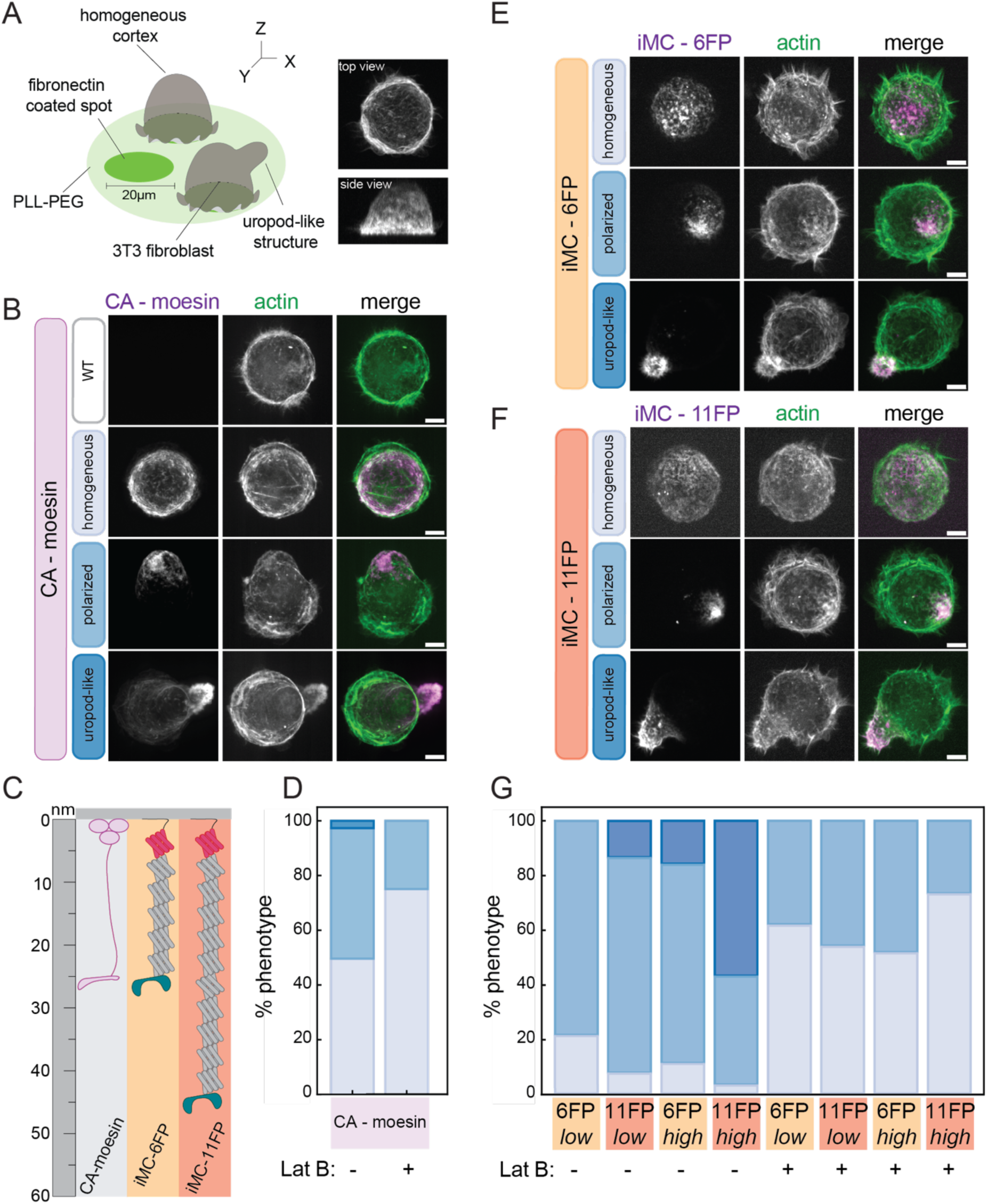
MCA linker protein caging regulates cortical mechanics in cells. A) Schematic of cells on micropatterns with representative top and lateral maximum intensity projections of dome-shaped NIH3T3 fibroblast cells on 20 μm micropatterns. B) Representative linker distributions of CA-Moe in NIH3T3 fibroblasts. C) Schematic of CA-Moe and engineered iMC-linkers used to assess the role of linker length. D) Quantification of the proportion of each phenotype in CA-Moe expressing cells with and without 500 nM latrunculin B (−LatB = 186 cells, +LatB = 145 cells, N=3). E-F) Representative linker distributions of iMC-6FP (E) and iMC-11FP (F) in NIH3T3 fibroblasts. G) Quantification of the proportion of each phenotype in high and low expressing clones of +iMC-6FP and +iMC-11FP in the presence and absence of the actin depolymerizing drug latrunculin B (N = 3 experiments, 6FP-Low = 156 cells, 6FP-High = 169 cells, 11FP-Low= 158 cells, 11FP-High= 160 cells, 6FP-Low +LatB = 147 cells, 6FP-High + LatB =159 cells, 11FP-Low + LatB = 122 cells, 11FP-High + LatB = 112 cells). See also Figure S6 and S7.

Our *in vitro* and *in silico* work show that membrane viscosity and linker length alter cortical mechanics by regulating the degree of linker caging. Thus, we examined the effects of manipulating both these parameters in cells. The classical method for perturbing membrane viscosity in living cells is via manipulation of membrane cholesterol content. Unfortunately, such interventions have also been shown to alter actin levels in the cell and disrupt important signalling pathways^80,81^. Nonetheless, this being the gold standard in the field we used this method to increase membrane viscosity in NIH3T3 cells by depleting cholesterol with 5 mM MβCD. As an alternative, we also increased membrane viscosity by decreasing the saturation levels of the plasma membrane by inhibition of the desaturase SCD1 with 2 mM A939572 + BSA 18:0^82^. Both treatments led to the expected changes in membrane lipid composition as assayed by lipidomics of whole cell fractions and plasma membrane isolates (**Figure S6A, B**). Moreover, both treatments lead to a reduction in the diffusion coefficient of a lipidated GFP (GPF-CAAX) (**Figure S6C-E**), with a concomitant cell rounding (**Figure S6F**). To decipher the mechanical determinants of the cell rounding phenotype we measured cell stiffness by AFM-nanoindentation on micropatterned cells (to rule out the effect of cell shape on the measurements) (**Figure S6G**). In agreement with the observed cell rounding, cortical stiffness increased in cells treated with A939572 or MβCD (**Figure S6H**). However, we also observed a reduction in the amount of actin in MβCD-treated cells (**Figure S6I, J**). This highlights the many effects of lipid manipulations and makes it difficult to interpret our findings for the understanding of the role of membrane viscosity on linker caging in cells.

We thus turned our focus to MCA linker length. To be able to distinguish between moesin’s signaling and tethering functions, we extended our repertoire of engineered artificial MC-linkers of varied lengths, to include ones which mimic the domain architecture of endogenous MCA proteins, but are inert with regard to signalling^74,83^. Specifically, they contain the CH1/CH2 domain of utrophin^84^ to bind F-actin, the lipidation motif of lyn^85^ to insert into the plasma membrane, and different numbers of fluorescent proteins to link the two domains and visualize their subcellular localization. We used a linker with a predicted length of over 30 nm (iMC-6FP-linker^74^), similar to active ezrin (25 nm^86^) and generated a novel one with a predicted length of over 50 nm (iMC-11FP-linker) (**Figure 6C**). For each construct, we generated doxycycline-inducible NIH3T3 fibroblast cell lines and isolated both high and a low expression clones (**Figure S7A**). The expression of such constructs in cells on micropatterns was sufficient to induce clear morphological changes, leading to the aggregation of actin and linker proteins into distinct caps, or even the formation of “uropod-like” structures in a manner that depended on linker length and density (**Figure 6E-G**). We called these structures “uropod-like” because they are morphologically reminiscent of the contractile tail that forms at the back of migrating immune cells^87^. To quantify this phenotype, we classified the linker distributions as homogeneous, polarized, or uropod-like. For low expression of the shorter linker, iMC-6FP-linker, we observed polarization of almost 80% of the cells. As linker expression and length increased, so did the proportion of uropod-like cells, with high expression of iMC-11FP-linker resulting in almost 50% uropod-like cells and only 4% of the cells remaining unpolarized. No cells had a phenotype with more than one uropod-like protrusion, suggesting that the wavelength of the pattern was on the order of the length of a cell or greater. To verify that the formation of tails resulted from the caging of MCA linker proteins within the actomyosin network, we depolymerized the cortex by adding 500 nM of latrunculin B. This treatment made all tails disappear, and also homogenized the distribution of linker proteins in cells (**Figure 6G, S7B**). Together, these results suggest the existence of a positive feedback between actin recruitment and caged linkers, whereby MCA linkers recruit actin, which in turn cages linkers. These results demonstrate that linker caging provides a powerful mechanism for the control of cortical mechanics and symmetry breaking in cells.

## DISCUSSION

We demonstrate a mechanism for the control of cortical mechanics and symmetry breaking in mammalian cells based purely on the mechanics and physical properties of the cell surface composite interface. This mechanism results from the caging of membrane-to-cortex attachment proteins within the underlying actin network. We first identify the relevant biophysical properties of the cell surface composite by using a minimal *in vitro* approach. By coupling a lipid bilayer via MCA proteins to an actin polymer network that is crosslinked and contracted by myosin motors, we show that membrane viscosity and linker length are tunable dials which together alter the ability of myosin to remodel the minimal actin cortex. Simulations and analysis of linker diffusivity indicate a mechanism for these phenomena: linkers become “caged” within the actin cortex. In cells, this mechanism leads to symmetry breaking, which depends on interactions between linker proteins and the actomyosin network. Expression of constitutively active moesin was sufficient to lead to symmetry breaking in NIH3T3 fibroblasts. This did not rely on the signaling properties of moesin: expression of artificial linkers without any signaling capacity had the same effect. They also confirmed the results obtained in the *in vitro* system by showing that the proportion and degree of these symmetry breaking phenotypes increased with linker length and expression level. Together, these results suggest that modulation of the biophysical parameters of the cell surface composite can be used by cells to control cortical dynamics and symmetry breaking.

### Linker caging: A powerful and tunable mechanism for controlling cell surface mechanics

Caging limits the ability of linker proteins to explore new filaments and regions by restricting their lateral mobility. Moreover, it constrains the mechanics and dynamics of the actin cortex, and leads to the formation of mechanically distinct domains. Most research on cortical heterogeneities and their effects on membrane patterning focuses on the regulation of myosin activity^88^. A prevalently used framework is the “picket fence model”. First proposed in the 1970s this model suggests that the cortical actin meshwork proximal to the surface acts as a “fence” anchored to the membrane via transmembrane proteins or “pickets”, which limit the movement of both lipids and proteins through steric hinderance or frictional effects. This temporary confinement (or corralling) creates the experimentally observed small membrane compartments with restrictive diffusion, which can act as regulators of neuronal polarization and receptor clustering in phagocytosis^89–91^.

In such a model, the membrane has always been viewed as a passive participant. Based on our results, we propose an active contribution of lipids and membrane proteins towards the formation of cortical heterogeneities. In our study, we explore two physical parameters – membrane viscosity and MCA protein length – that cells could use to locally tune cortical actin dynamics. It should be noted, however, that the control of linker caging occupies a complex multiparametric space. Membrane composition, via alterations of solvation properties, electrostatic potential, charged density and lipid packing^57^, can further contribute to the regulation of caging through interactions between structural features of linker proteins and membranes. In fact, while our simplified *in vitro* system removed many of these variables in order to isolate the effects of membrane viscosity and MCA protein length, even here even here we observed that M-moe and S-moe interact differently with cholesterol-containing membranes in a manner which cannot be accounted for by length alone. Specifically, even though our supported lipid bilayers (SLB) are completely fluid (**Figure S1A**), M-moe fluorescence did not fully recover on cholesterol-containing membranes in FRAP experiments (without MAC, **Figure S5B**). S-moe fluorescence, in contrast, recovered fully (**Figure S5A**). We hypothesize that this is due to the presence of additional alpha-helix and acidic flop domains in the M-moe construct. These domains are highly enriched in tandem repeats of positively-charged Lysine residues and thus increase the possibility of electrostatic interactions with membranes (as has been previously shown^92,93^). Furthermore, properties of the actin cortex, such as density, pore size, filament length, and cortical architecture^94^, as well as the membrane-to-cortex distance,^75^ will also modulate the ‘effective’ binding of the linker proteins (**Figure 5B-C, S8E**). Linker structure and affinity for both the membrane and for actin likely play important roles (**Figure S8A, S8E**). Indeed, we can find some evidence for this when comparing our data to the literature. The expression of constitutively active ezrin, a paralogue of moesin with which it shares extensive structural (**Figure S8C**) and sequence conservation^95^, leads to symmetry breaking in T-cells^36^ and melanoma cells^96^ but fails to do so in fibroblasts (**Figure 6A-D** *vs.* **S8B**). We attribute the failure of ezrin to break symmetry in fibroblasts to the lower binding affinity of the ezrin actin binding domain when compared to that of moesin (**Figure S8A**). However, that it does break symmetry in some cells suggests that different cell types can exploit the full multiparametric space to regulate cell surface mechanics in a context-dependent manner. Adding another layer of complexity, membrane and cortical features not only individually regulate aspects of the caging interaction, but also exhibit complex interdependencies. For example, the charge of membranes is expected to influence membrane-to-cortex distance^73^.

Finally, while our work has focused on the cell surface, composite interfaces are also present at the periphery of the nucleus^97^ and other membranous organelles, as well as at membrane contact sites^98^, and similar phenomena may well be active at these sites. Each of these distinct subcellular environments presents unique mechanical requirements and constraints. Thus, to fully understand the regulatory potential of living composite interfaces, tools and techniques will need to be developed to independently and simultaneously study multiple parameters with precision in cells.

### Membrane viscosity and MCA protein length are emergent regulators of cytoskeletal dynamics at the cell surface

By combining minimal *in vitro* systems with MCA linker protein engineering, we reveal two physical parameters – membrane viscosity and MCA protein length – that cells employ to tune cortical actin dynamics.

As our experiments to isolate the role of membrane viscosity in linker caging demonstrate, great care is needed to interpret the consequences of manipulations of lipid composition. This is due to the myriad signaling effects of lipids and the current technical impossibility of manipulating lipid mechanical properties without simultaneously changing cellular signaling. Despite technical limitations, however, an appreciation is beginning to emerge for the immense diversity of lipid types and the importance of their precise regulation^99^, not only for signaling purposes, but also for the maintenance of the mechanical integrity of the cell. In fact, cells actively regulate membrane viscosity via changes in lipid composition, most commonly through the regulation of the ratio of cholesterol and sphingomyelin and that between saturated *vs*. unsaturated lipids in the membrane^13,14^. The regulation of membrane fluidity in response to external stresses such as temperature^14^ or mechanical stress^100,101^ is well characterized. Additionally, lipid composition and membrane fluidity are intrinsically regulated according to cell type, state, and cell cycle stage^102–105^. Despite great progress, a comprehensive understanding of the significance and implications of lipid diversity is still lacking^13^. We propose that the role of membrane properties in the regulation of the cell surface composite should be included in future considerations. Targeted alterations of plasma membrane fluidity could be employed by cells to regulate how effectively myosin can remodel the cortex. As our *in vitro* results show, small changes in membrane composition can be amplified to create large effects on cell mechanics. Interestingly, a two-fold change in membrane fluidity, such as the one in our *in vitro* experiments, has been estimated between subcellular compartments^106^ or in diverse dynamic states^21^.

Information is transmitted from the plasma membrane to the actomyosin cortex via MCA proteins. Murrell and Gardel observed a decrease in the magnitude of actomyosin network strain when increasing the binding to the underlaying membrane^59,71^, in line with MCA proteins mediating adhesion between the membrane and the cortex. In addition to density, we identify MCA linker length as another critical parameter. Long linkers get caged withing the cortex, functionally entangling actin filaments and limiting myosin mediated remodeling. The concept of the physical size of proteins acting as an important determinant of their distribution in cells has interesting precedents in the literature. Chang and Dickinson, for example, report that the size of PAR-3 oligomers determines their sorting in the developing C. elegans embryo, a direct result of size-dependent interactions with the actomyosin cortex^107^.

Our results thus raise the question as to how great the diversity of MCA protein lengths is in cells. The ezrin-radixin-moesin (ERM) protein family is the first and by far most completely characterized family of MCA proteins. The impressive conservation of ERM proteins, homologues of which have been identified across all metazoans^108^, provides an important clue as to their fundamental importance in cellular mechanisms that have persisted throughout evolution^95,109^. Although there are no studies of ERM protein structure *in situ*, it is reasonable to hypothesize that lengths will be similar, based on their high sequence homology (see **Figure S8C** for Alpha fold prediction). Early studies on ezrin from placental microvilli suggest the presence of large quantities of noncovalent ezrin oligomers^110^. While detailed structural information on these oligomers is lacking, dimerization or multimerization of ERM proteins provides a possible mechanism by which cells might modulate linker length. Moreover, while ERM proteins are the best studied family of MCA proteins, other families of proteins that simultaneously bind the plasma membrane and actin will also effectively act as linkers. This includes flotillin (discussed further below), BAR domain containing proteins like Snx33^74^, talin, or unconventional myosin motors^111^. Additionally, the association with scaffolding proteins, such as between ezrin and EBP50^112^, might increase effective linker length and thus affect the “cageability” of these proteins.

Our results show that changing membrane viscosity and linker length can not only result in complete stalling of cortical networks, but also result in changes in timescales, as the 2x slowdown in cortical rearrangements seen in **Figure 4B** for S-moe. An understanding of how this subtler mechanism of regulation is used in cells will require more precise biomolecular tools to probe time-dependent processes. However, it already provides a mechanism which synthetic biologists could exploit in their attempts to reproduce processes such as cell division in synthetic cells. Perhaps synthetic biology efforts could in turn feed new hypothesis as to the use of this mechanism in cells.

### Symmetry breaking at the cell surface

The cell surface is an immensely complicated system, and spatial heterogeneity on a microscale is key to its function. Processes such as endo and exocytosis, for example, require temporary accumulations of proteins to bend the membrane^113^, and numerous signaling processes exploit the assembly of domains enriched in specific protein and lipid species^114–116^. The cell must maintain a delicate balance: it must permit the heterogeneities necessary for cell function while containing the positive feedback loops that would drive it into undesired asymmetry.

The search for general principles which underlie and enable this delicate balance has often focused on the amplification mechanisms and feedback loops by which small signals or fluctuations can be transformed into cell-wide phenomena^1^. This focus harkens back to the theoretical work of Turing who proposed a model for biological pattern formation based on reaction-diffusion equations^117^. Later studies based on this idea have since offered many insights into how patterns are generated and spread across cells^118^. We observe only a single uropod-like or ‘cap’ structure per cell, suggesting that the wavelength of the perturbation is on the length scale of a cell. A better understanding and mathematical description of our system will reveal whether and how cells can functionally tune this wavelength.

Much work on symmetry breaking has focused on the feedback loops that control the polymerization, structure, and contractility of the actomyosin cytoskeleton via Rho/Rac/CDC42^11^. Others, informed by a physics perspective, have focused on the buildup of forces and tension in the unstable, active cellular system^119–121^. Such mechanisms place a strong focus on the role of the actomyosin cortex, but the membrane, and in particular membrane tension, has also been shown to be a global inhibitor of actin polymerization, regulating symmetry breaking in migrating cells^122^. Together, these different mechanisms are beginning to build up a picture of a cell poised for symmetry breaking via the presence of multiple systems on the brink of instability. Here, we propose that linker caging is a fundamental underlying control mechanism which cells use as a regulatory dial to control the establishment and maintenance of symmetry breaking events.

ERM proteins are canonical polarity markers for a range of symmetry breaking phenomena^27^. We will take the formation of the uropod, a characteristic “tail-like” structure found at the back of cells undergoing amoeboid migration^87^, as a case study to show how the mechanism of linker caging synergizes with and potentially explains previous experimental observations: to initiate the formation of a uropod in polarizing immune cells, ezrin is first globally deactivated by dephosphorylation. This suggests that the presence of active ezrin, perhaps caged in the cell cortex, has a stabilizing effect, and that removing this inhibitor then creates a permissive environment for cell polarization. Subsequently, a smaller subset of active ezrin is localized to the ‘cap’ domain which initiates the formation of the uropod. Formation of this cap is dependent on F-actin, but independent of myosin-driven actin contractility^36,37,123^. FRAP experiments of constitutively active ezrin in the cap suggested that the mobility of ezrin is significantly restricted when active. Treatment with latrunculin A abolished the difference between constitutively active and wild type ezrin, indicating that interaction with F-actin restricts the mobility of the active form^36^. Similarly, the lipid raft scaffolding protein flotillin has also been proposed to function in membrane-to-cortex attachment during uropod formation^123^. It was shown that deletion of either the membrane-binding or the actin-binding domain of flotillin-2 abolishes its ability to localize to the cap, and latrunculin treatment increased the mobility of flotillin-2 in T cells stimulated to induce uropod formation^123^. Together, these studies support a role for protein caging in the establishment and maintenance of polarity, and specifically for the integrity of the back of migrating cells.

Another process for which there is strong evidence for a role for linker caging in symmetry breaking is the generation of apicobasal polarity in the developing mouse embryo. Ezrin localizes to the apical surface of cells during embryonic polarization at the 8-cell stage^33^. An exploration of this process suggested that actin network density and protein mobility in the membrane were important for the successful aggregation of ezrin clusters and symmetry breaking, and that disrupting the balance between these two biophysical parameters impaired symmetry breaking. In particular, the authors suggest that a higher density actin network aids the cooperative assembly of ezrin clusters, and disruption of actin with Jasplankinolide or ARP2/3 inhibitors resulted in a failure of symmetry breaking. In summary, linker caging appears to be a fundamental mechanism underlying cell polarization, and this notion provides a conceptual framework with predictions to be explored in the future.

We propose that the composite nature of the cell surface provides emergent biophysical dials that the cell can exploit to regulate symmetry breaking phenomena. Our *in vitro* experiments allowed us to identify membrane viscosity and MCA linker length as novel regulators of cortical mechanics. An important task faced by the cell is how to spatially restrict and regulate processes involving interactions at the cell surface at physiologically relevant timescales. We propose that the biophysics of the living composite interface at the cell surface allow the cell to restrain the dynamics of this system, maintain stability, and break symmetry when required by cellular processes.

### Limitations of the study

While our MACs provide fundamental mechanistic insights, they do not recapitulate the complexity of living systems in terms of linker diversity, membrane composition and heterogeneity, as well as actin turnover, crosslinking, branching and length distributions. These additional factors very likely contribute to when, where, and under what conditions linker caging occurs in cells. Moreover, we monitor lipid/protein dynamics at an ensemble level using FRAP. However, this could miss important effects that emerge from small heterogeneities in lipid or protein distributions.

Our coarse-grained simulation models aim at identifying the minimal physical ingredients that give rise to the observed phenomena. As such they implicitly represent the membrane by a homogeneous viscous medium, they model linkers as floppy polymers and neglect hydrodynamics. In particular, the presence of hinges or mechanical resistance as linkers fold might in principle affect the system dynamics.

The artificial linkers that we use in cells allow us to assess the biophysical roles of linker proteins but differ in important ways from endogenous proteins, such as their binding affinity for actin (**Figure S8 A**), and likely their flexibility. Our linkers are also always active, and are thus lacking the on/off kinetics of normal linker proteins. Additionally, while we have included AlphaFold predictions of the sizes of the different constructs in the manuscript, these are estimates and the precise length of our linkers and the conformation they adopt in cells or in our SLB system may differ from these estimations. Finally, the pleiotropic effects of lipid manipulations in living cells make it complicated to tease apart their effects on linker caging.

## ACKNOWLEDGMENTS

We thank Thomas Pucadyil, Anna Erzberger, Thomas Quail and Christina Kurzthaler for critical reading of the manuscript and useful discussions. We thank the EMBL flow cytometry core facility, and the EMBL advanced light microscopy facility for support and advice. Also, Qin Yu and Anna Kreshuk for help with image analysis.

## AUTHOR CONTRIBUTION

S.D, M.L and A.D-M conceived the project. S.D, R.T.M, A.B.G, C.S.E and A.D-M designed the experiments. S.D. performed the *in vitro* experiments. R.T.M and A.B.G. performed the *in cellula* experiments. R.R.S and A.B.G performed the lipidomics experiments under the supervision of C.S.E and A.D-M. Z.G.S analyzed the dynamics of actin remodeling *in vitro* under the supervision of M.M. I.P developed the computational models and performed and analyzed computer simulations of MCA linkers of various lengths/viscosities under the supervision of A.Š. J.B discussed simulations. S.D, S.K.F and A.D-M wrote the manuscript. All authors contributed to the interpretation of the data, read, edited and approved the final manuscript.

## SUPPLEMENTAL VIDEOS

**Video S1.** FRAP on Texas Red-labelled free DOPC, Chol and DOPE membranes. Related to Figures 1 and 2. Scale bar = 5 µm. t = sec.

**Video S2.** Actomyosin dynamics on DOPC membranes with MAC. Actin is labelled with Phalloidin Alexa-488. Scale bar = 10 µm. t = min:sec.

**Video S3.** Actomyosin dynamics on Chol membranes with MAC. Actin is labelled with Phalloidin Alexa-488. Scale bar = 10 µm. t = min:sec.

**Video S4.** Actomyosin dynamics on DOPE membranes with MAC. Actin is labelled with Phalloidin Alexa-488. Scale bar = 10 µm. t = min:sec.

**Video S5.** Simulations of actomyosin dynamics with variable fluidity.

**Video S6.** Simulations of actomyosin dynamics with small (S), medium (M) and long (L) linkers.

**Video S7.** Simulations of caging with small (S), medium (M) and long (L) linker.

## MATERIALS AND METHODS

### Protein purification

#### Moesin constructs

Residues 1-577 of human moesin were cloned with a C-terminal StrepII tag in a pET-15B vector with a phosphomimetic T558D mutation^124,125^ (L-moe). Residues 1-301 and 1-501 were deleted to generate M-moe and S-moe constructs, respectively. N-terminal GFP-tag moesin (L-moe: 1-577 aa, and M-moe 302-577 aa) T558D, were custom designed and ordered from Genewiz in the pET-15B and pRSET vector (His-TEV-GFP-thrombin-moesin-Strep-Tactin®) respectively and S-moe (302-577 aa) was subsequently subcloned from GFP-L-moe. All clones were confirmed by Sanger sequencing. Proteins were expressed in BL21(DE3) competent cells (PepCore, EMBL) in terrific broth supplemented with 1.5% lactose, 0.05% glucose, and 2 mM MgS0_4_ at 18°C for 18 or 30 hours (non-GFP and GFP-tagged proteins, respectively) and purified in a 2-step purification. For constructs in pRSET vectors, bacterial pellets were immediately processed. For proteins in pET-15B backbone, frozen bacterial pellets were resuspended in 20 mM Tris, 500 mM KCl, and 10 mM imidazole pH 7.4 (filtered and degassed), supplemented with a protease inhibitor cocktail (Roche, 5892791001) and subsequently lysed in a microfluidizer (Microfluidics-M-110L; 3 cycles). Proteins were first purified and eluted with a HiTrap IMAC HP column (Cytiva, 17092003) against 150 mM imidazole, followed by binding to a StrepTrap HP column (Cytiva, 29401320) and eluted against 2.5 mM disthiobiotin. For storage, proteins were dialysed against Tris-KCl (20 mM Tris, 150 mM KCl) supplemented with 5 mM *β*-mercaptoethanol, 1 mM EDTA, and 10% v/v glycerol with final pH of 7.4 overnight, flash frozen in liquid N_2_ and stored at −80 °C for 3 months. Prior to experiments, proteins were dialyzed overnight in Tris-KCl buffer (20 mM Tris, 150 mM KCl, pH 7.4) and spun at 100,000×g for 20 minutes to remove glycerol and aggregates.

#### Actin and skeletal muscle myosin

Actin (AKL99-A) and skeletal muscle myosin (MY02-A) were purchased from Cytoskeleton Inc., reconstituted as per instructions, and snap frozen in experiment-sized aliquots. Briefly, before an experiment, actin was thawed, incubated with G-buffer (20 mM Tris, 0.2 mM CaCl_2_, 0.2 mM ATP and 0.5 mM DTT, pH 8.0) overnight, and then centrifuged at 60000 rpm for 30 minutes at 4°C in a Beckman TLA 100.3 rotor to remove any aggregates. The concentration of G-actin in the supernatant was determined on a NanoDrop 1000 Spectrophotometer (ThermoScientific) at A_290nm_ using *ε* (2.66*10^4^ M^-1^cm^-1^). Catalytically dead myosin motors were removed as in^58^. The concentration of active myosin after centrifugation was determined on a spectrophotometer at A_280nm_ using *ε* (2.638*10^5^ M^-1^cm^-1^).

### Preparation of fluid supported lipid bilayers (SLBs)

p-Texas red DHPE was isolated from Texas Red DHPE (Invitrogen, T1395MP) using silica gel plates as in^126^. All lipids were purchased from Avanti Polar Lipids: DOPC (850375C), DOPS (840035C), DGS-NTA(Ni^2+^) (790404C), Chol (700000P), DOPE (850725C), DOPIP_2_ (850155P). Lipid stocks were reconstituted in chloroform in the following proportions to generate Chol-, DOPE- or PIP_2_ plus NTA– containing mixtures: DOPC:DOPS:DGS-NTA(Ni^2+^):TxR (82.5:12:5:0.5 mol%), DOPC:DOPS:DGS-NTA(Ni^2+^):Chol:TxR (52.5:12:5:30:0.5 mol%), DOPC:DOPS:DGS-NTA(Ni^2+^):DOPE:TxR (22.5:12:5:60:0.5 mol%) and DOPC:DOPS:DGS-NTA(Ni^2+^):DOPIP_2_:TxR (77.5:12:5:5:0.5 mol%). Stocks were stored at −20°C and brought to RT before use. Supported lipid bilayers (SLBs) were generated by spreading 1.5 μL of the lipid stock in the center of a glass coverslip which had been chemically treated and passivated with PEG8000 as in^126^.

### Actomyosin network assembly for contractility experiments

To generate minimal actin cortices (MACs), 5 μM of F-actin was first polymerized in KMEI Buffer (20 mM Tris, 100 mM KCl, 0.2 mM EGTA, 1 mM MgCl_2_, 0.2 mM ATP, 1 mM DTT) for 90 minutes at RT and then placed on ice to stop the reaction. In a typical experiment, SLBs were generated inside the flow-cell in KMEI buffer. Once a uniform SLB formed, 300 µL of unlabeled moesin (unless specified otherwise) was flowed in and incubated for 10 minutes followed by 3 washes with 1 mL each of KMEI buffer to remove unbound protein. Longer incubation times interfered with the stability of the SLB (**Figure S2A**). Next, a 2 µM solution of F-actin was briefly incubated with 0.5 µM of Alexa488 Phalloidin (Thermo Fisher, A12379). The combined solution was gently pipetted with blunt-cut 200 μL pipette tips and immediately flowed into the chamber. Following 10 minutes of incubation with the SLB, unbound actin was washed out. Next, a 100 nM solution of myosin was incubated in KMEI buffer (final volume of 300 µL) at RT for 10 minutes to allow formation of minifilaments before being flowed in to the chamber. For all imaging experiments KMEI buffer was supplemented with an oxygen scavenger cocktail of 0.2 mg/mL glucose oxidase (Sigma, G-2133), 0.035 mg/mL catalase (Sigma, C-40), and 4.5 mg/mL glucose. For all the reactions with myosin, the KMEI buffer was supplemented with 1 mM of ATP regenerating mix (from a 1X stock of 10 mM Mg-ATP, 100 mM creatine phosphate, and 100 U/mL creatine kinase). All reactions were carried out at 25°C.

### Electron Microscopy

Electron microscopy on myosin II filaments was performed using minifilaments at 0.01 μM as in^127^.

### *In vitro* fluorescence microscopy

Fluorescence imaging was performed on a Nikon Ti-E inverted microscope equipped with a CFI P-Apo DM 60x Lambda oil-immersion objective with a numerical aperture (NA) of 1.40. A Lumencor Spectra X light engine was used as an illumination source and emission was collected through single-band filters with excitation/emission wavelengths of 485 ± 25 nm/525 ± 50 nm for Alexa488, 560 ± 2 nm/593 ± 40 nm for Texas red probes, and 650 ± 13 nm/692 ± 40 nm for Alexa647 probes using a pco.edge 4.2 sCMOS camera (Excelitas Technologies). Image acquisition was controlled by NIS Elements Viewer (Nikon).

### Fluorescence Recovery after Photobleaching (FRAP)

FRAP was done on the same system using identical software for imaging and analysis for *in vitro* and cell experiments. Imaging was carried out on a Zeiss LSM 780NLO laser scanning confocal microscope by photobleaching a region of interest. The focus was set to the bottom plane of the cell or to the membrane patch (in case of *in vitro* experiments) with maximum fluorescence, and a circular region of >1 µm was bleached. Image acquisition was controlled by the Zen software (Zeiss). Imaging parameters were standardized to minimize photobleaching during acquisition. For every bleached ROI, a reference ROI and a background ROI were chosen manually and the signal intensity in each of the three regions was estimated for each frame of the movie. All further analysis was performed using the FRAPAnalyser software^128^. First, the intensity in the bleached region was background-subtracted and further normalized against the reference region at each timepoint. For 2D diffusion in the membrane and considering a circular bleached region, the intensity recovery over time was modeled as in^129^:

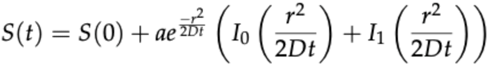

where *S* is signal intensity, *t* is the time after bleaching, *r* is the radius of the bleached spot and *I*_0_ and *I*_1_ are modified Bessel functions arising due to the circular geometry of the bleached region. The factor *a* accounts for incomplete recovery of the signal and can be expressed as *S*(∞) − *S*(0). The above equation was fit to the averaged FRAP curve to obtain fitted values of the diffusion coefficient, *D*. Averaged recovery curves from all recorded data per condition were plotted using the Matplotlib package on Python 3.6.1 and box plots of fitted D values per cell were plotted using Tableau Desktop 2020.4. for data in cells and using Graphpad Prism (version 10.0) for data in *in vitro*.

### Image Analysis of *in vitro* experiments

#### Filament length

Measurements of myosin II minifilament and actin filament lengths were performed manually in ImageJ (version 2.9.0/1.52u).

#### Quantification of protein binding

For quantification of GFP-labelled S-, M-, and L-moe constructs and Alexa 488-Phalloidin labelled actin, 100X100 pixel ROIs in the membrane channel were used as reference to select and crop ROIs in F-actin or moesin channels from an 2044X2060 pixel image (1-4 independent ROIs/image). Background for the two channels was extracted from ROIs (1-4 ROIs/image) lacking membrane for every experiment. After background-correction, mean intensity was extracted from all the pixels in the ROI in the protein channels. A Gaussian filter (0.5, median) was applied to every image for representation purpose in manuscript. All imaging analysis was done using ImageJ (version 2.9.0/1.52u) and all statistical analysis was carried out using Graphpad Prism (version 10.0).

#### Laurdan Imaging and Analysis

For Laurdan-based experiments all lipid mixes (DOPC, +Chol and +DOPE) were supplemented with 0.5:0.5 mol% of rhodamine (Avanti, 18:1 Liss Rhod PE, 810150P) and C-Laurdan (bio-techne, #7273). Imaging was carried out on a Zeiss LSM 780NLO laser scanning 2-photon confocal microscope using a 63x Oil DIC M27 objective with NA of 1.4. The dye was sequentially excited with the 561 nm (for the rhodamine channel) and 2-photon (800 nm) laser lines and emission was read at 568-630 nm for rhodamine, 414-457 nm (for the liquid order-Lo phase), and 491-535 nm (for the liquid disorder-Ld phase). The Generalized Potential (GP) analysis was performed in ImageJ (version 2.9.0/1.53u), according to the following equation:

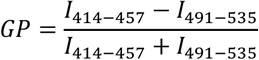

where *I* is the intensity of pixels in Lo and Ld phases. GP values were calculated for each pixel of a membrane patch (250X250 pixels) of a 16bit image manually using the image calculator function in ImageJ.

#### Nematic order calculations

First, the background of image data was subtracted using ImageJ’s Background Subtraction function with a window size of 50 pixels. The nematic order parameter, <N>, was calculated using a customized MATLAB script (versions 2019b and 2023a, Mathworks) as in^58^. Briefly, the local F-actin orientation was determined for each direction field of a small windows of 19x19 pixels with 50% adjacent overlaps. Each window was Gaussian filtered and transformed into Fourier space using a 2D fast Fourier Transform (FFT). The axis of the least second moment was determined from the second-order central moments of the transformed window, and the angle of the local F-actin orientation was defined as orthogonal to this axis. Next, the local degree of alignment was calculated between adjacent windows within 3×3 kernels. We then calculated the local nematic order for the central window in each kernel using the modified order parameter equation:

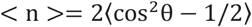

where θ is the difference in F-actin orientation between the central window and the 8 surrounding windows. We repeated this process for all possible 3×3 kernels over an image, yielding a nematic director field with defined director magnitude and orientation for each window over an image. Perfect alignment between adjacent regions within an F-actin network results in an order parameter of one.

#### Particle image velocimetry (PIV)

Particle image velocimetry (PIV) was applied to the fluorescent actin images using a customized MATLAB script (version 2019b and 2023a, Mathworks) as in^58^. The extent of contraction was calculated by defining a strain 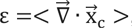, as the divergence of the displacement field. The data was analyzed with PIV window size 32 and overlap 0.5. Background subtraction was conducted in ImageJ on the F-actin channel (50-pixel window size) to enhance the signal-to-noise ratio (SNR).

#### Calculation of bundling parameter

We performed the filament bundle analysis using the 2D image thresholding method. Epifluorescence images were first background subtracted in ImageJ, using a 50-pixel radius. Images were then thresholded in MATLAB using a customized script. Briefly, we rescaled the image by dividing each pixel by the sum of all pixel intensity values in the image. All the values ≤ 0 were then excluded, leaving only high-intensity pixels likely belonging to filaments and bundles. Lastly, we calculated the mean and standard deviation of the remaining regions, and used those to obtain the bundling parameter λ:

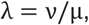

where ν is the standard deviation, and µ the mean.

### Simulations of contractile actin networks

#### Overview

To understand how linker-actin binding affects actin contractility (**Figure 3A-B, 4E-F**, **Videos S5, S6**), we developed a coarse-grained computational model. Actin filaments were represented as semirigid polymers; myosin was represented by a two-bead dumbbell, each end of which was able to bind an actin bead and directionally walk toward the barbed end through a bond flip; linkers were represented by floppy polymers anchored to a fluid surface from one end and binding to actin filaments at the other end. All the three components were composed of spherical volume-excluding beads connected by springs. Myosin attachment and detachment, myosin walking, and linker binding and unbinding were stochastic and occurred at set rates. Simulations were run with three different initial actin architectures; for each initial architecture all linker lengths and viscosities were tested.

Simulations proceeded in two phases: an equilibration phase followed by a production run. In the equilibration phase, linker-actin binding was allowed but myosin-actin binding was forbidden mimicking our *in vitro* experiments before myosin addition. In the production run, myosin-actin binding was switched on, making the actin network connected and contractile. Simulations were performed by overdamped molecular dynamics in LAMMPS^130^ with the use of the REACTER package^131^ and results are visualized in OVITO^132^.

#### Actin, myosin, and linkers

Each actin filament was modelled as a polymer of 100 spherical beads, connected by a harmonic bond of equilibrium length σ and harmonic constant *k*_stretch_ = 500 kT, where *k* is the Boltzmann constant and *T* is temperature, and by a harmonic angle potential of strength *k*_bend_ = 500 kT that keeps the polymer straight. Actin filaments were therefore unstretchable and fairly rigid, with a persistence length 10 times larger than the filament length. Each linker was modelled as a polymer of spherical beads of diameter σ (that we took as the simulation length unit), connected by a harmonic bond of equilibrium length σ and harmonic constant *k*_stretch_ = 500 kT. Linkers were therefore unstretchable and floppy with lengths 4, 9 and 20 σ (respectively, S, M and L in **Figures 4E-F**). Each myosin unit was modelled as a dumbbell of 2 beads, connected by a spring of equilibrium length 10 σ and harmonic constant *k*_stretch_ = 20 kT. Volume exclusion between beads was modeled through a Weeks-Chandler-Anderson (WCA) potential of amplitude ɛ_repulsion_ = 10 *kT* and range *σ*.

#### Time integration

The dynamics resulted from a velocity-Verlet integrator of the equations of motion, coupled with a Langevin thermostat, such that the simulated ensemble was NVT. The Langevin thermostat adds a viscous drag force and a random force to the force originating from inter-particle interactions, thus implicitly representing collisions with the solvent. The viscous drag force is −𝑚𝒗/𝜏_th_, where 𝒗 is the velocity of the particle and 𝜏_th_ is the characteristic damping time of the thermostat; 𝜏_th_ can therefore be interpreted as an inverse viscosity. For **Figures 4E-F** and **Video S6**, the characteristic damping time of the thermostat was set to 𝜏_th_ = 0.1 𝜏, with 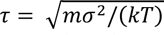 the simulation time unit and 𝑚 the mass of a bead. For **Figures 3A-B** and **Video S5**, we set 𝜏_th_ to 0.1 𝜏, 0.05 𝜏 and 0.001 𝜏, to obtain a 2-fold and 100-fold increase in medium viscosity that mimicked changes in membrane viscosity. Such small 𝜏_th_ values ensured that the system was in the strongly overdamped regime, where inertia is negligible, and favoured numerical stability despite the continuous energy exchanges due to myosin dynamics (see below).

An equilibration phase lasting 2 · 10^4^ 𝜏, during which myosin was inactive, was followed by a production run lasting 20 · 10^4^ 𝜏. The time steps, chosen to avoid instabilities, were 𝜏_𝑠_ = 0.005 𝜏 for **Figures 4E-F** and **Video S6**, and 𝜏_𝑠_ = 0.001 𝜏 for **Figures 3A-B** and **Video S5**.

#### Simulation box and densities

The first bead of each linker was constrained to move on the plane 𝑧 = 0, representing the membrane. The remaining particles initially occupied the region 3 σ ≲ 𝑧 ≲ 20 σ (actin) and 0 ≲ 𝑧 ≲ 20 σ (linkers and myosin), but they were compressed during equilibration to the region 3 σ ≲ 𝑧 ≲ 6 σ (actin) and 0 ≲ 𝑧 ≲ 6 σ (linkers and myosin). This allowed us to reach higher actin densities and ensured that the vast majority of all actin filaments were bound by at least one linker throughout all our simulations. These spatial constraints were enforced by Lennard-Jones walls of range 0.5 σ and repulsion strength 10 𝑘𝑇. We varied the number of actin filaments between 100 and 600, in a simulation box of side 𝐿_𝑥_ = 𝐿_𝑦_ = 200 σ, which corresponds to a post-equilibration actin packing fraction between 4 and 26%; we show data for 400 filaments in **Figures 3A-B** and **Video S5** and 600 filaments in **Figures 4E-F** and **Video S6**. The number of linkers was 4000. The number of myosin motors was 400, low enough for the actomyosin network to be dynamic and not jammed without linkers.

#### Linker and myosin binding dynamics

From the beginning of the simulation, we allowed linker tips to bind to unbound actin beads at a rate of 1 𝜏^−1^ for any two such beads that are in contact. The formed bond is harmonic, with equilibrium length 1.2 σ and strength 20 𝑘𝑇. Linker unbinding (bond deletion) occurred at a rate of 10^−3^, 10^−4^ or 10^−5^ 𝜏^−1^ (we show results for 10^−4^ 𝜏^−1^ in **Figures 3A-B** and **Video S5** and for 10^−5^ 𝜏^−1^ in **Figures 4E-F** and **Video S6**); results were qualitatively similar, but we observed that the actin dynamic arrest is more pronounced for longer lasting bonds, consistent with our following observation that the increase of caging time with linker length becomes more pronounced for stronger bonds (**Figure S8E**). Myosin beads could bind unbound actin beads with the same rate and bond parameters mentioned for linkers. A myosin bead could walk along an actin filament by moving the bond that connects it to an actin bead to the next actin bead, in the direction of the barbed end. This occurred with rate 10 𝜏^−1^, provided that the next actin bead was not already bound to another myosin and was within distance 1.5 𝜎. We assumed that the myosin walking rate was unaffected by linkers possibly bound along the filament. Myosin unbound from actin (bond deletion) at a rate 10^−4^ 𝜏^−1^. The myosin unbinding dynamics did not depend on the position along the actin filament, resulting in myosin dwelling at barbed ends.

The presented data result from 3 simulations with different random number generator seeds (hence different realizations of the initial architecture and of the dynamics).

#### Data analysis

The actin mean square displacement (**Figure 4F**) was determined by averaging the squared displacement since the beginning of the production run over all actin beads.

### Simulations of linker dynamics

#### Overview

To understand how linker-actin binding affects linker dynamics (**Figure 5A-C, S8D-F, Videos S7**), we developed a coarse-grained computational model where linkers were represented by polymers anchored to a fluid surface at one end and binding to steady actin filaments at the other end. Both linkers and actin filaments were composed of spherical beads connected by springs. All beads interacted *via* volume exclusion; the last bead of the linker interacted with actin beads *via* an additional short-range attraction. The system was simulated through overdamped molecular dynamics in LAMMPS^130^ and results were visualized in OVITO^132^.

#### Setup

Each linker was modelled as a polymer of spherical beads, connected by a harmonic bond of equilibrium length σ and harmonic constant 𝑘_stretch_ = 1000 kT. Linkers were therefore unstretchable and floppy. Linker lengths were varied between 6 and 48 σ (in **Figure 5A-C** we label 6 σ as S, 9 σ as M, 24 σ as L). The first bead of the linker was constrained to move on the plane 𝑧 = 0. A Lennard-Jones wall positioned at 𝑧 = −0.5 σ, of range 0.5 σ and repulsion strength 10 𝑘𝑇 kept all particles in the 𝑧 ≳ 0 region. The cortex occupied the region 5 σ < 𝑧 < 50 σ and was composed of immobile actin filaments. The filament orientation and size was random in the following sense: upon initialization of the system, a random point in the cortex region and a random unit vector were chosen as the origin and direction of a new filament. Beads were placed at distance σ in the direction of the unit vector until either the filament reached the maximum length of 50 beads, or the last bead happened to be at the boundary of the cortex region or overlapped with a previously placed filament. The filament was then ended and the procedure repeated for a new filament. In **Figure 5A-C** the total amount of actin was limited to 5000 beads, for a box of length 𝐿_𝑥_ = 𝐿_𝑦_ = 100 σ, yielding a relatively low packing fraction of ∼ 1%. In **Figure S8D** this system was compared to a denser one, with 25000 beads and ∼ 5% packing fraction.

Volume exclusion between beads was modeled through a Weeks-Chandler-Anderson (WCA) potential of amplitude ɛ_repulsion_ = 10 𝑘𝑇 and range 𝜎, for both linker-linker and linker-actin interactions. After a pre-equilibration (see below), we allowed linkers to bind actin. This was done by setting the interaction between the last bead of each linker and any actin monomer to the sum of a WCA and a cosine-squared potential: namely, the force was repulsive for distances < 0.8 𝜎 as per WCA and attractive for distances between 0.8 𝜎 and 1 𝜎. We varied the characteristic energy of this interaction, ɛ, between 6 and 14 𝑘𝑇 (**Figure S8E**; **Figure 5B-C** presents results for 8 𝑘𝑇).

#### Time integration and viscosity

The dynamics resulted from a velocity-Verlet integrator of the equations of motion, coupled with a Langevin thermostat, such that the simulated ensemble is NVT (see **Simulations of contractile actin networks** for the thermostat action). The chosen time step was 𝜏_𝑠_ = 0.01 𝜏, where 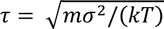 was the simulation time unit and 𝑚 the mass of a bead. The characteristic damping time of the thermostat 𝜏_th_, which can be interpreted as an inverse viscosity, was to 1 𝜏 in the cortex region; it was varied between 0.1 𝜏 and 10 𝜏 in the region below the cortex, which represented the membrane and the space between membrane and cortex, to mimic changes in membrane viscosity. These values are small enough to ensure that the system is in the overdamped regime for time scales above 10 𝜏 = 10^3^𝜏_𝑠_.

The system was pre-equilibrated for 5 ⋅ 10^6^ 𝜏_𝑠_, during which linkers, initially oriented vertically, relaxed to their compact (folded) state. Attractive interaction with actin was then switched on and the system was further equilibrated for another 5 ⋅ 10^6^ 𝜏_𝑠_. After that, a 1 ⋅ 10^8^ 𝜏_𝑠_ production run started. The presented results are averages over the 100 linkers present in each simulation, and over *N* simulations with equal parameters and different random number generator seeds (hence different realizations of the cortex and of the dynamics): *N* = 8 for **Figure 5C and S8D-F**, *N* = 3 for **Figure 5B**.

#### Data analysis

From the simulations we computed linker mean square displacement projected on the *xy* plane (**Figure S8F**): the dynamics were diffusive at low times, caged at intermediate times, and again diffusive at long times. We extracted the caging time, i.e. the typical time at which linkers recovered diffusion after being trapped in the cortex, and the long-time diffusivity. The caging time grew and the diffusivity decreased with linker length at a rate that cannot be explained by the corresponding increase in linker mass which scales linearly with length. As expected, caging time also increases and diffusivity decreases with bond energy ɛ (**Figure S8E**), showing that these phenomena depend on binding dynamics. More interestingly, the relative increase in caging time and decrease in diffusivity with linker length become more pronounced for stronger bonds. In addition, caging increases and diffusivity decreases when actin packing fraction is increased from 1% to 5%, i.e. when the cortex meshes are smaller (**Figure S8D**). Altogether, this suggests that longer linkers get trapped in the cortex meshes, because after unbinding they tend to rebind to filaments within the same cortex mesh, before their tips can diffuse away from the mesh. This massively slows down their dynamics.

### Cell culture

NIH 3T3 fibroblasts (CRL-1658, ATCC repository) were cultured in Dulbecco’s Modified Eagle Medium (DMEM) with 4.5 g/L D-glucose, supplemented with 10% fetal bovine serum (Thermo Fisher, 10270-106, lot 2384425) and 1% penicillin-streptomycin. Cells were cultured at 37°C with 5% CO_2_ on polystyrene dishes. Cells were passaged every 2-3 days (up to 70% confluency) using 0.05% trypsin-EDTA. Cells were routinely tested for mycoplasma contamination.

### Cloning

The iMC-6FP-linker and CA-Ezrin constructs were cloned as described in^74^. The iMC-11FP-linker was generated by introducing five dark mCherry (Y72S) into the iMC-6FP-linker using endonuclease cloning into a PiggyBac vector (kindly provided by Jamie Hackett’s group, EMBL Rome). The phosphomimetic human moesin was taken from the pET15B vector (*in vitro* cloning) into a PiggyBac vector for transduction of cells. All plasmids were confirmed by Oxford nanopore sequencing.

### Cell line generation

All the cell lines expressing synthetic linkers were generated using a PiggyBac transposon system that includes a doxycycline(dox)-responsive TRE3G promoter. Cell lines were generated by co-transfecting the PiggyBac vector and the PiggyBac transposase via lipofectamine 3000 (company). Transfected cells were selected using 1.2 mg/mL of geneticin in the culture medium. Single clones were obtained by performing serial dilutions of cells into 96 well plates, and screened for fluorescence levels via flow cytometry (Flow Cytometer BD Symphony A3 Analyzer). Clones with low fluorescence background and similar expression levels upon dox induction were selected (**Supplementary Figure S7A**). 16 hours prior to any experiment linker expression was induced by the addition of 1 µg/mL dox in the medium.

Lentiviral transduction was used to generate CAAX-GFP 3T3 fibroblasts. Briefly, lentivirus particles carrying the GFP-CAAX gene under a constitutive promoter were produced in HEK293T cells. These particles were collected from the HEK293T supernatant and used to transduce 3T3 cells, which were subsequently sorted by fluorescence-activated cell sorting (FACS) to obtain a population of cells with a homogeneous level of stable GFP-CAAX expression.

### Cellular micropatterning on glass

NIH3T3 fibroblasts were seeded on 20 µm adhesive spots on glass dishes. Adhesive spots were generated using an inverted Ti2^®^ Nikon microscope equipped with the Primo micropatterning module (Alvéole). In brief, 35 mm glass-bottom dishes (Greiner bio-one, 627860) were plasma treated for 1 minute (Gala Instrument, Plasma Prep 2) and a small PDMS stencil with an inner hole of ∼10-20 µL was placed in the middle of the dish to reduce reagent usage. Next, the glass surface inside the stencil was coated with a non-adhesive layer of 0.2 mg/mL polyethylene glycol (PLLg-PEG, Susos AG) for 1 hour at RT. Then, the surface was washed three times with PBS before applying a thin layer of 4-benzoylbenzyl-trimethylammonium chloride micropatterning reagent (PLPP, Alvéole). Using a 375 nm laser (4.5 mW) 20 µm circles spaced 40 µm apart were photo-patterned onto the glass surface, using the µmanager software v.1.4.22 with the Leonardo plugin v.4.12 (Alvéole). Finally, the dish was coated with 50 µg/mL bovine fibronectin (Merk, F1141) for 30 minutes and washed 3 times with PBS. After washed, 10^5^ cells were seeded onto the patterned dish. Cells were allowed to settle and adhere for 20 minutes before non-attached cells were washed out. Experiments were performed 2 hours after cell seeding if not indicated otherwise.

### Drug treatments

To modify lipid composition, cells were treated with either A939572 (Merck, SML2356) to inhibit SCD1, or Methyl -β - cyclodextrin (MβCD, Merck, C4555) to deplete cholesterol. For this, lyophilized A939572 was dissolved in DMSO to a stock concentration of 4 mM and stored at −20°C. Stearic acid (18:0) was complexed with 3 mM BSA in 3T3 medium as described for fatty acid supplementation, to a final stock concentration of 6 mM. To perform SCD1 inhibition in cells, the A939572 stock was dissolved in 3T3 media to obtain a final concentration of 2 µM. Media from the 3T3 cells in culture was aspirated and replaced with the 2 µM A939572 solution. 4 hours later, the BSA-18:0 stock was added to the cells at a 1:30 dilution, in order to have final concentrations of 200 µM 18:0 and 100 µM BSA. The cells were then co-incubated with A939572 and BSA-18:0 over 16-20 hours to achieve increased saturation of their lipids as validated by mass spectrometry (**Figure S6B**).

Cholesterol was depleted using 5 mM MβCD. In order to prevent competition between cellular cholesterol and lipids in the media for binding MβCD, the drug was dissolved in cell media made with delipidated FBS (7BioScience, 7-FBS-DL-12B) in place of regular FBS. Cells were washed twice with PBS before being placed in media containing the drug to wash out any lipid remnants that could interfere with the binding of MβCD to cellular cholesterol. Cells were incubated with the drug for 30 minutes before collection for lipidomic analysis (**Figure S6A**). For patterned cell imaging 30-45 minutes of incubation time was used. To depolymerize the cell cortex, cells were treated with 500 nM of latrunculin B (Merck, 428020) for 15 to 45 min. The drugs were kept in the media during image acquisition.

### Fluorescence microscopy in live cells

Unless otherwise specified, fluorescence imaging was acquired on an Olympus -IXplore SpinSR microscope equipped with a UPLSAPO100XS Silicon objective with an NA of 1.35. The cells were imaged by acquiring z-stacks of 0.28 µm with a 2x ORCA-FLASH4.0 V3 camera. Cells were imaged live at 37°C and 5% CO_2_ after being seeded onto micropatterns 2-4 hours before imaging. Prior to image acquisition, the cells were incubated with 1:1000 SPY650-FastAct (SC505, Spirochrome) diluted in DMSO (as per manufacturer’s instructions) for 1 hour. As negative staining for cell segmentation AF 488 NH-ester (11820, Lumiprobe, prepared as per manufacturer’s instructions) was added 1:1500 to the media during image acquisition.

NIH3T3 fibroblasts expressing CA-moesin were imaged on a Nikon Ti-2CSU W1 SORA Spinning Disk microscope with a SR P-Apochromat IR AC 60x water immersion objective with an NA of 1.27. The cells were imaged by acquiring z-stacks of 0.30 µm with a Hamamatsu OrcaFusionBT camera. For image acquisition, the CS-W1 SoRa mode was utilized to increase magnification.

### Image Analysis in cells

#### Actin mean fluorescence intensity

z-stacks of cells were acquired in 3 channels: our expressed constructs, a live actin reporter and a negative stain.

For 3d segmentation, we trained a convolution neural network using pytorch-3dunet^133^ and plant-seg^133^ with manually curated actin segmentations obtained in ilastik^134^ and the negative channel as the segmentation channel. Next, the negative channel was used to generate masks for surface quantification. Actin mean fluorescence intensity (**Figure S6J**), was then calculated by applying the masks across the z-stacks. The mean actin fluorescence of each cell was normalized to the mean intensity of control cells with no cholesterol depletion.

#### Symmetry breaking analysis

Phenotypes were classified by analyzing the distribution of the linkers at the cell surface. Maximum intensity projections of each cell were used as an overview for manual classification. If the phenotype was not clear, the individual z stack for the cell was further inspected to distinguish across phenotypes. The phenotypes were classified as ‘homogeneous’, if there was a continuous distribution in the cortex; ‘polarized’, if the linker formed a cap within the cell cortex; and ‘uropod-like’, if the linker accumulation led to a change in topography in the cell as observed in the representative images (**Figure 6B, 6E** and **6F**).

### Single-cell atomic force spectroscopy

Nano-indentation was performed on a CellHesion® 200 atomic force spectrometer (Bruker) integrated into an Eclipse Ti® inverted light microscope (Nikon). Measurements were run at 37 °C with 5% CO2 and samples were used for no longer than 1 hour for data acquisition. Measurements were acquired at a sampling rate of 2 kHz and analyzed using the JPK Data Processing Software.

Nano-indentation was performed using tipless MLCT-O10 type C cantilevers (spring constant ∼0.01 N/m; Bruker), calibrated using the thermal noise method. A silica bead with a diameter of 10 μm (Microparticles) was glued onto the cantilever using a 2-part epoxy resin (UHU) with 5 minutes working time. Upon full hardening of the glue, the cantilever was coated for 30 minutes at 37 ° C with a 1% (w/v) Pluronic F-68 (Thermo Fisher Scientific) in milli-Q water and rinsed with PBS, aiming to prevent non-specific adhesion to the probed cells. Force-distance curves were acquired using 500 pN contact force and 0.4 μm/s approach/retract velocity. Up to 5 curves were taken per cell, with a 10 s waiting time between successive curves to prevent history effects.

Cortical tension (T_c_) was determined by fitting each force-indentation curve between 0 (the contact point) and 300 nm with a liquid droplet model suited for nano-indentation experiments^135^:

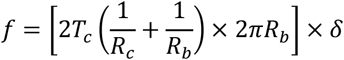

Where *f* is the force applied to the cell surface which leads to a displacement (δ) of cellular material; *R_c_* and *R_b_* are the radius of the cell and the silica bead glued on the cantilever, respectively.

### Cell lipidomics

#### Whole cells

3*10^6^ ± 0.5*10^6^ NIH3T3 fibroblasts were collected for whole-cell lipidomics. For this, they were trypsinized, counted, and pelleted. The cell pellet was washed once with 4 ml HKM buffer (150 mM HEPES-KOH, 150 mM KCl and 5 mM MgCl2, pH 7.4), re-pelleted and washed once more with 1 ml HKM buffer. The final cell pellet was then dissolved in 200 µl of 155 mM ammonium formate, transferred to a 2 ml Eppendorf tube, and snap-frozen in liquid nitrogen for storage at −80°C until further processing.

#### Plasma membrane isolation

Plasma membranes were isolated by differential centrifugation using the Minute Plasma Membrane Protein Isolation and Cell Fractionation Kit (Invent Biotechnologies, SM-005), as per the instructions of the manufacturer. Briefly, NIH3T3 fibroblasts were seeded at a density of 5*10^3^ cells/cm^2^ on four 15 cm dishes. Three days later, the cells were trypsinized and pelleted by centrifuging at 500-600×g for 5 minutes at RT. The cells were washed once with PBS at 4°C, counted, and centrifuged again at 500-600×g for 5 minutes. Typically, 25*10^6^ ± 10*10^6^ cells were retrieved in this manner. The supernatant was aspirated completely and the cell pellets were all combined by resuspending in 500 µl of buffer A and incubated on ice for 5-10 minutes, following which they were vortexed for 10-30 seconds. Samples at this stage are termed whole cell lysate (WCL). The WCL was then transferred immediately to a filter cartridge provided in the kit, capped, and centrifuged at 16000×g for 30 seconds at 4°C. The pellet thus obtained was resuspended by vigorous vortexing for 10 seconds and passed through the same filter once again by repeating the centrifugation step. Following this, the filter was discarded and the pellet was resuspended once again by vigorous vortexing for 10 s. The sample was centrifuged at 700×g for one minute. The supernatant was transferred into a fresh 1.5 ml Eppendorf tube and centrifuged at 16000×g for 30 minutes at 4°C. The supernatant was carefully removed and the pellet was resuspended in 200 µl of buffer B by vigorous vortexing for 30 seconds and centrifuged at 7800×g for 5 minutes at 4°C. The supernatant was carefully transferred to a fresh 2 ml Eppendorf tube and mixed with 1.6 ml PBS at 4°C by inverting the tube a few times. This mixture was then centrifuged at 16000×g for 30 minutes at 4°C. The supernatant was discarded and the pellet containing the isolated and washed plasma membrane fraction (PM) was resuspended in 200 µl of 155 mM ammonium formate.

Protein concentration in isolated plasma membranes was estimated using Bradford or BCA assay and the samples were diluted to a final protein concentration of 45 ng/µl in 155 mM ammonium formate, following which they were snap-frozen in liquid nitrogen for storage at −80°C until further processing.

#### Lipid extraction

Lipids were extracted from the samples as described previously^136^. Briefly, all samples were thawed at 4°C and homogenized by sonication, following which they were spiked with an internal standard mix prepared in-house as described in^137^. Lipids were extracted using a two-step method^138^, first by partitioning the sample against chloroform/methanol (10:1, vol/vol) and then using the aqueous fraction to again partition against chloroform/methanol (2:1, vol/vol). The solutions were shaken using a ThermoMixer (Eppendorf) set at 1400 rpm and 4°C during both steps (for 2 and 1.5 h, respectively) and the organic phases were collected. Solvents were then evaporated under vacuum, leaving deposits of lipids extracted in the 10:1- and 2:1-partitioning steps, respectively. The 10:1- and 2:1-extracts were dissolved in chloroform/methanol (1:2, vol/vol).

Shotgun MS^ALL^ lipidomic analysis was performed as previously described^136^. Briefly, 10:1- extracts were diluted with 2-propanol to yield an infusate composed of chloroform/methanol/2-propanol (1:2:4, vol/vol/vol) and 7.5 mM ammonium formate for positive ion mode analysis and similarly with chloroform/methanol/2-propanol (1:2:4, vol/vol/vol) and 0.75 mM ammonium formate for negative ion mode analysis. The 2:1-extracts were diluted with methanol to yield an infusate composed of chloroform/methanol (1:5, vol/vol) and 0.005% methylamine for analysis in the negative ion mode. These were injected into an Orbitrap Fusion Tribrid mass spectrometer (Thermo Fisher Scientific) using a robotic nanoflow ion source, TriVersa NanoMate (Advion Biosciences). High-resolution mass spectra of intact ions (MS1) and fragments generated from 1 Da windows of intact ions (MS2) were recorded using an Orbitrap mass analyzer. The m/z peaks of intact and fragmented ions of lipids were assigned using ALEX123^139,140^ and the molar abundance of lipid molecules was quantified by comparing their MS1 peak intensities to that of spiked-in internal lipid standards. Lipid quantification and analysis of fatty acid composition were carried out on the SAS9.4 platform (SAS) as described previously^141^.

### Statistical analysis

Statistical analyses are described in all figure legends and were performed using R or GraphPad. Data visualization was done using both R or GraphPad. Normality of data distribution was tested by Shapiro-Wilk test. A two-tailed t-test was used for normally distributed data when only 2 conditions had to be compared. Otherwise, a non-parametric Wilcox test was used. For multiple hypothesis testing one-ANOVA (Kruskal-Wallis multiple comparison test) was used. In all box-plots, the lower and upper hinges correspond to the first and third quartiles (the 25th and 75th percentiles). Black/solid line corresponds to the median. In violin-plots the colored area reflects the probability distribution of the data points.

---

**Figure S1.**
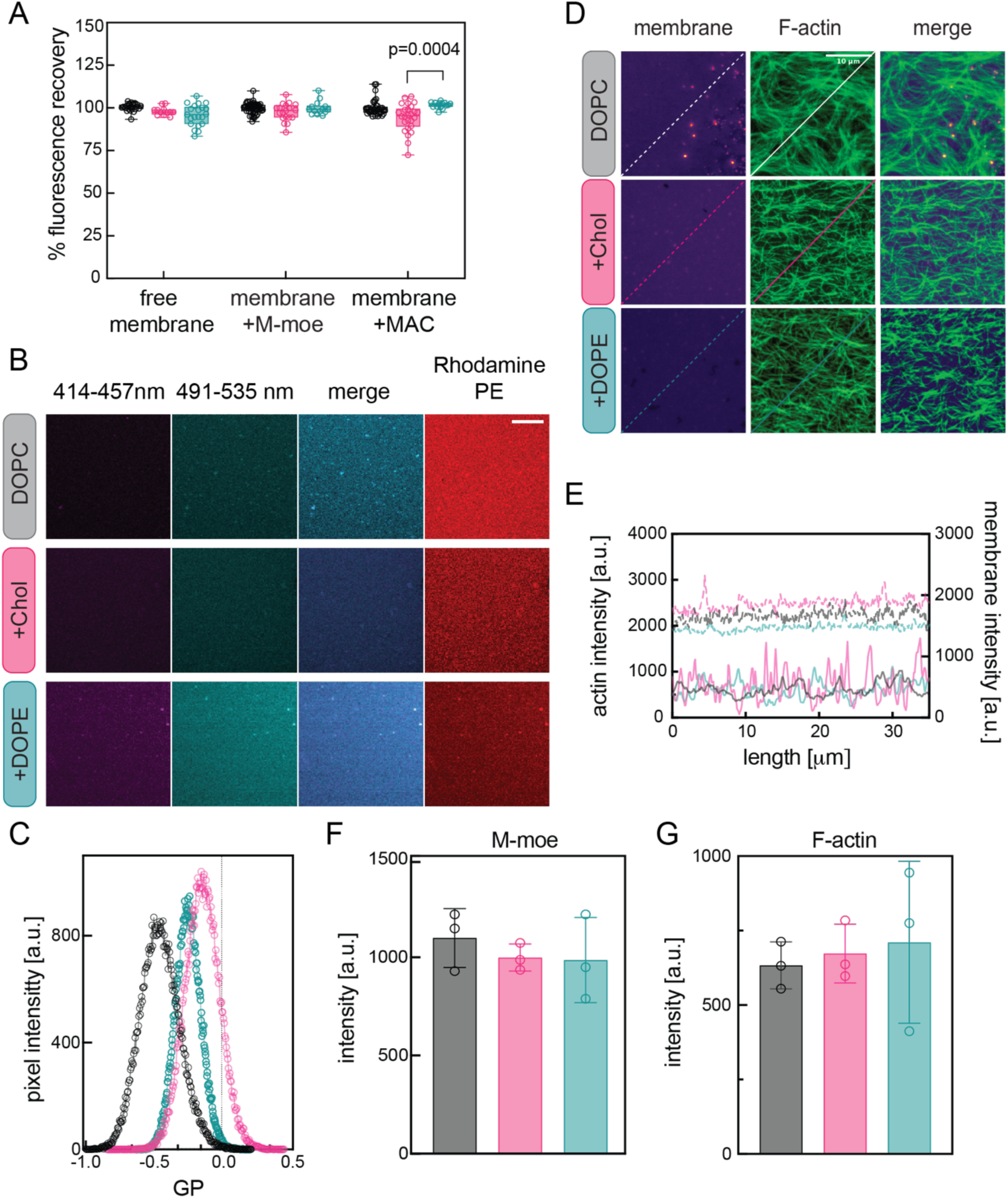
Characterization of MAC. A) Percentage recovery of Texas Red-labelled lipids on free membranes, membranes + M-moe, and membranes + MAC on DOPC (grey), +Chol (pink) and +DOPE (teal) membranes. B) Representative images of supported bilayer with C-Laurdan at 414-457 nm (for the liquid ordered-Lo phase, in magenta) and 491-535 nm (for the liquid disordered-Ld phase, in cyan) and merge for DOPC, +Chol, +DOPE membranes. The right panel shows bilayers in the RhodaminePE (0.5 mol%) channel to visualize membranes in Ld phase (no phase separation observed at room temperature). C) Histogram of Generalized Potential (GP) values of images in B (see **Methods** for details). D) Representative MAC images in the membrane, F-actin (labelled by Alexa 488-Phalloidin) and merge channels on DOPC (grey), +Chol (pink) and +DOPE (teal) membranes. E) Representative line profiles from (D). Solid lines indicate intensity distribution in the actin channel and dotted lines indicate intensity across the image in the membrane channel. F) Mean GFP-M-moe intensity bound to Texas Red-labelled membranes (DOPC (grey), +Chol (pink) and +DOPE(teal)) before actin addition (N=3,3,3 experiments). G) Mean intensity of Alexa 488-Phalloidin-labelled actin bound to Texas Red-labelled DOPC (grey), +Chol (pink) and +DOPE (teal) membranes via unlabelled M-moe (n>5 images/experiment, N=3,3,3 experiments). a.u. = arbitrary units. Solid line in box plots represents median. For significance: One-way ANOVA (Kruskal-Wallis multiple comparison test).

**Figure S2.**
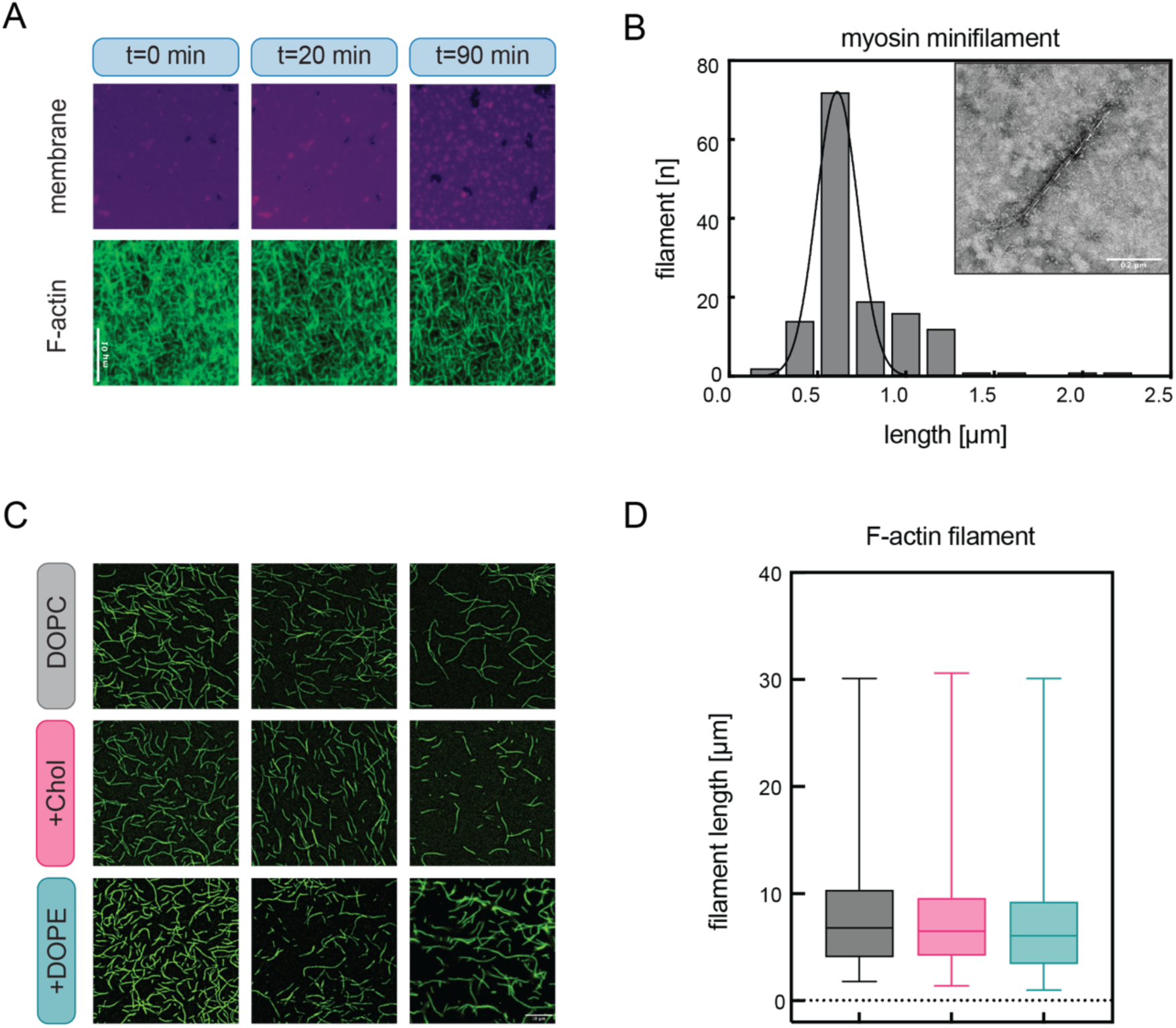
Characterization of actin and myosin. A) Representative time-lapse epifluorescence images of Alexa 488-Phalloidin-labelled actin network on Texas Red-labelled membranes. B) Distribution of myosin II filament length obtained from EM images at a concentration of 0.01 µM as shown in inset, white dotted line overlaying myosin minifilament indicates length as measured. C) 3 representative epifluorescence images (from left to right) of individual actin filaments stabilized with Alexa 488-Phalloidin bound to DOPC, +Chol and +DOPE membranes via unlabelled M-moe at concentration of 1 µM, 100 nM, 10 nM respectively. D) Box plots of actin filament length distributions (N=2 experiments) on DOPC (grey), +Chol (pink) and +DOPE (teal) membranes. Scalebars = 10 µm in A and C and 0.2 µm in B. Solid line in box plot represents median. For significance: One-way ANOVA (Kruskal-Wallis multiple comparison test), none of the comparisons were statistically significant.

**Figure S3.**
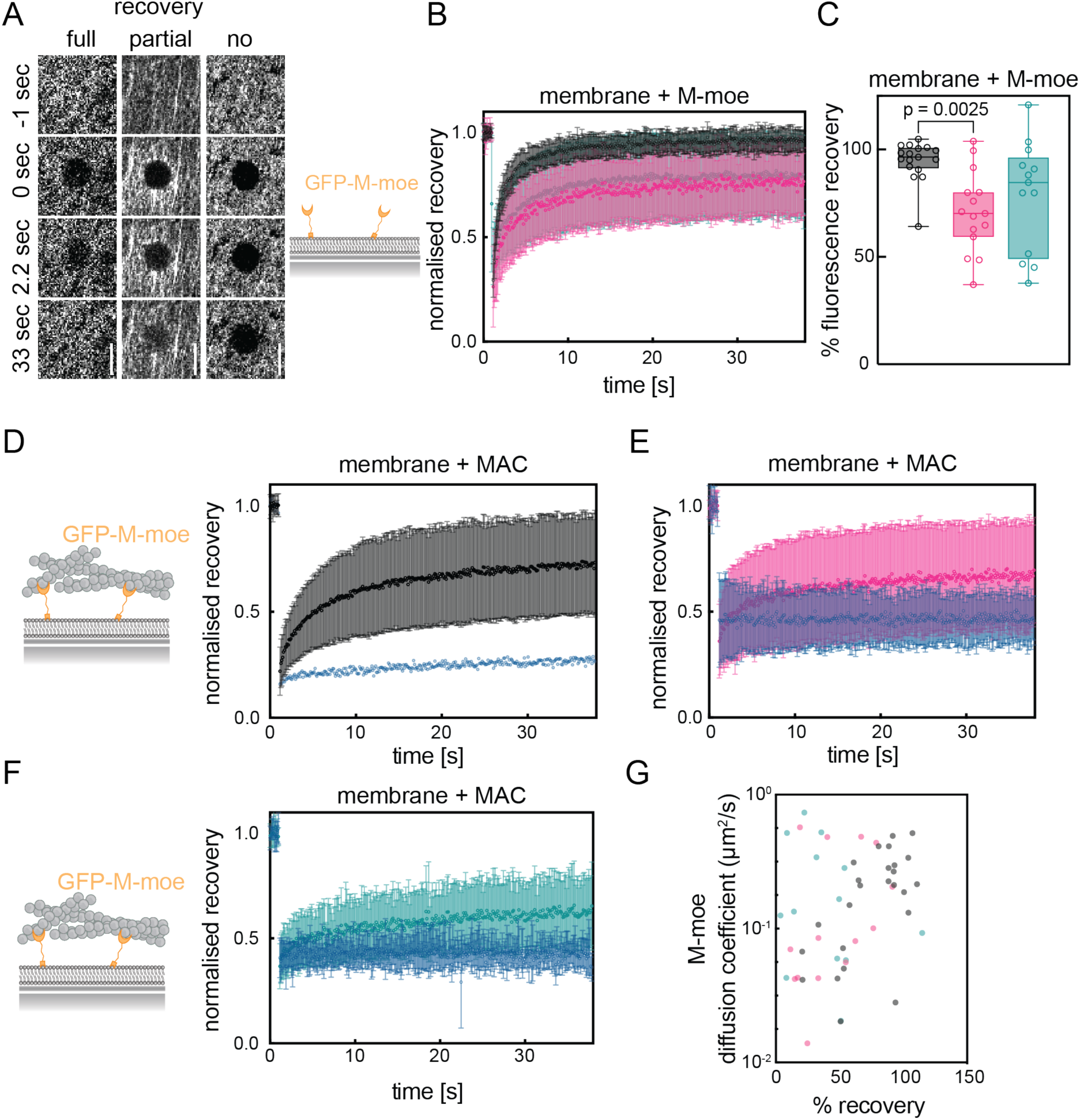
FRAP curves. A) Representative time-lapse images from independent FRAP experiments of GFP-M-moe with varying fluorescence recovery levels (see Table S1). B) Averaged traces from FRAP experiments of GFP-M-moe on DOPC (grey), +Chol (pink) and +DOPE (teal) free membranes. C) Percentage recovery of (B). D-F) Averaged traces from FRAP experiments of GFP-M-moe on DOPC (grey), +Chol (pink) and +DOPE (teal) membranes with MAC. Each trace shows an average of all curves: On each plot, light grey traces on (DOPC), pink (+Chol), teal (+DOPE) represent curves in which fluorescence recovered and blue traces represent curves in which no fluorescence recovery was observed (n≥3 curves/experiment, N≥3 experiments). G) Scatter plot of diffusion coefficient *vs.* percentage recovery of fluorescence of GFP-M-moe on DOPC (grey), +Chol (pink) and +DOPE (teal) membranes with MAC quantified from recovered curves only. Scale bars in A and B = 5 µm. Solid line in box plots represents median and error bars indicate SD. For significance: One-way ANOVA (Kruskal-Wallis multiple comparison test). See also Table S1.

**Figure S4.**
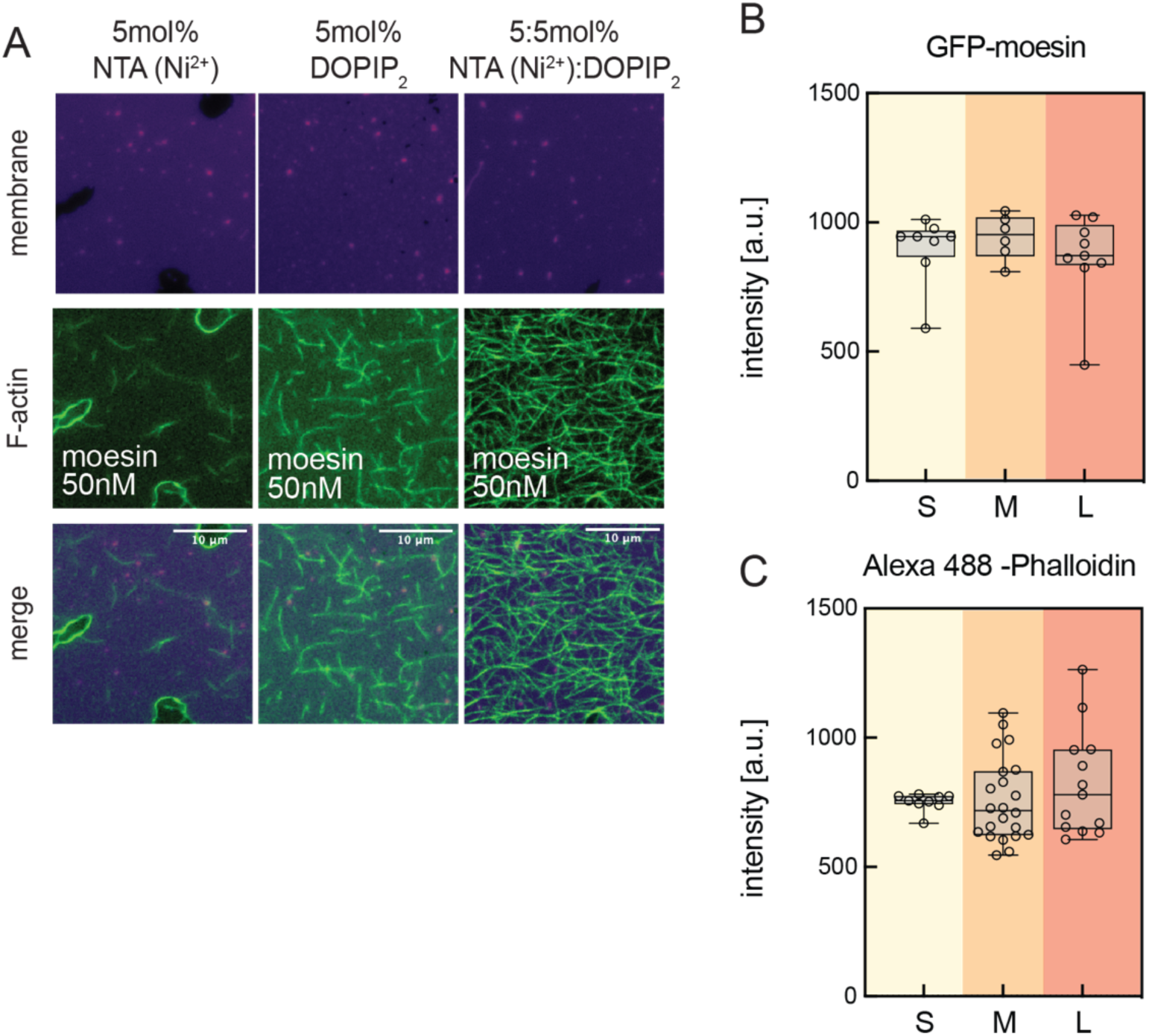
Characterization of MACs with S, M and L-moe constructs. A) Representative images in the membrane and actin channels acquired after actin recruitment on membranes doped with either 5 mol% NTA (Ni^2+^) or 5 mol% PIP_2_ or both coated with L-moe. Due to our short incubation time (10 min, see **Methods** for details) both NTA (Ni^2+^) and DOPIP_2_ were necessary for recruitment of the full-length protein. B, C) GFP-moesin and F-actin intensity distribution quantified on DOPC membranes coated with S-, M-, or L-moe (n≥2 images/experiment, N=3 experiments). Scale bars in A = 10 µm. a.u. = arbitrary units. Solid line in box plot represents median. For significance: One-way ANOVA (Kruskal-Wallis multiple comparison test).

**Figure S5.**
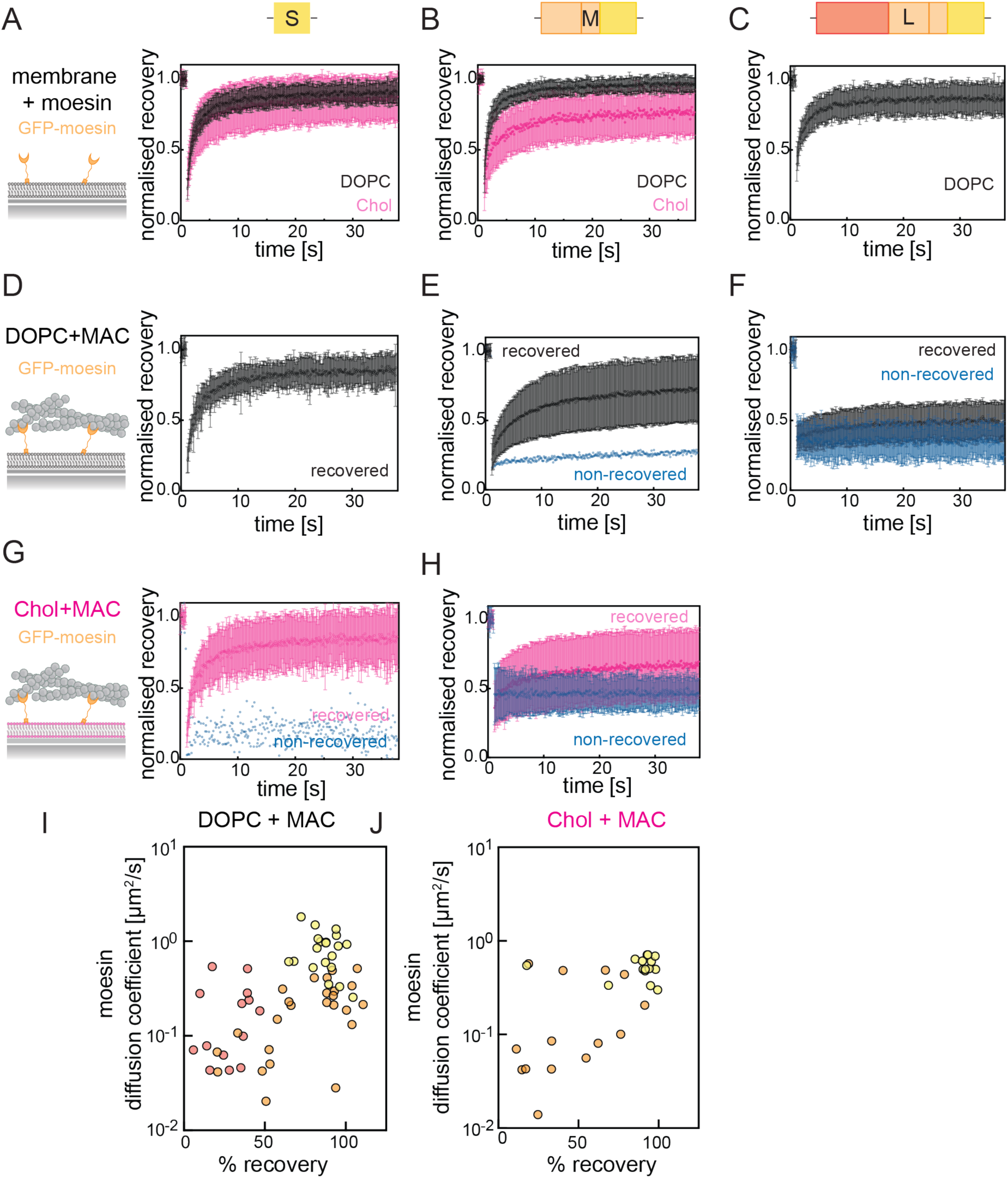
FRAP curves. A-C) Averaged traces from FRAP experiments of GFP-labelled S-, M-, and L-moe constructs on DOPC (black) +Chol (pink) membranes. Each trace shows an average of all curves. D-F) Averaged traces from FRAP experiments of GFP-labelled S-, M-, and L-moe constructs on DOPC membranes with MAC. On each plot, traces in black represent recovered curves and blue represent non-recovered curves. G-H) Averaged traces from FRAP experiments of GFP-labelled S-, M- moe constructs on +Chol membranes with MAC. On each plot, traces which show recovery of fluorescence are in pink and those which show no recovery are in blue. I-J) Scatter plot of diffusion coefficient *vs.* percentage recovery of GFP-moesin on DOPC (grey) (I) and +Chol (pink) (J)-membranes with MAC quantified from recovered curves only (4 <N>10 curves/experiment from at least N=3 experiments). See also Table S2.

**Figure S6.**
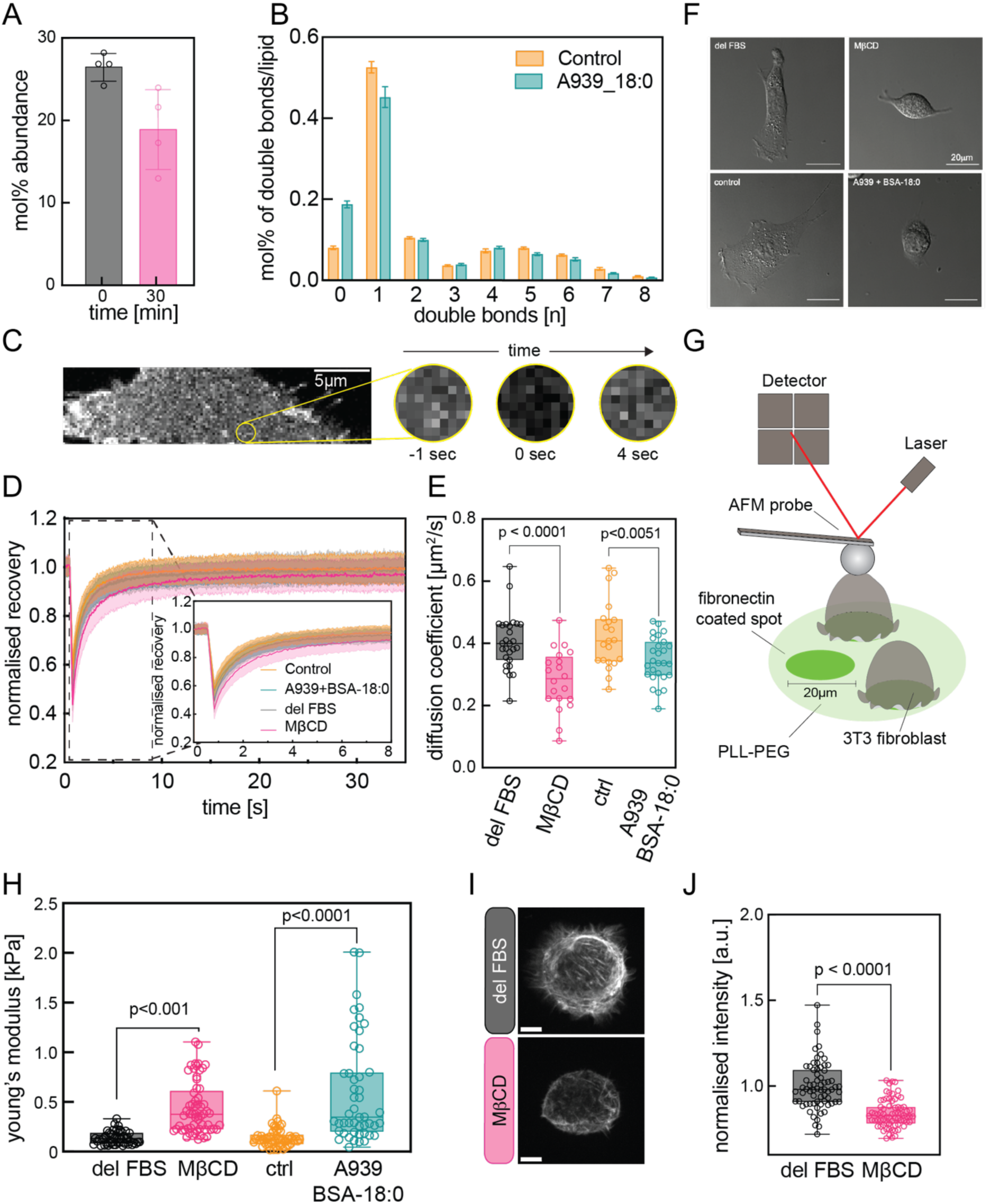
A decrease in membrane viscosity by cholesterol depletion or inhibition of the SCD1 desaturase leads to an increase in cortical stiffness. A) Mol% abundance of cholesterol in plasma membrane isolates upon 5 mM treatment with MβCD. B) Mol% of double bonds per lipid molecule before and after treatment with A939572 + BSA loaded with stearic acid (18:0). C) Representative confocal image of the basal membrane of a NIH3T3 fibroblast expressing the membrane marker CAAX-GFP before and after photobleaching. D) Recovery curves after photobleaching. E) Diffusion coefficient upon perturbation of lipid composition by MβCD or A939572 + BSA 18:0. F) Representative bright field images of NIH3T3 fibroblasts before and after addition of MβCD and A939572 + BSA 18:0 showing rounding up. G) Experimental setup for cortical stiffness measurements on micropatterned dishes by nano-indentation. H) Cortical stiffness measurements upon perturbation of lipid composition by MβCD or A939572. I-J) Representative images (I) and quantification of F-actin (J) in NIH3T3 fibroblasts in control and MβCD treated cells (N = 3 experiments, delipidated FBS (delFBS) = 67 cells and MβCD = 71 cells). Each dot represents one cell. In all other box plots dots represent the mean of multiple measurements of a single cell unless specified otherwise. Scale bars = 5 μm, a.u. = arbitrary units. Normality of data distribution was tested by Shapiro-Wilk test. Two-tailed t-test was used for normally distributed data. Otherwise, a non-parametric Wilcox test was used.

**Figure S7.**
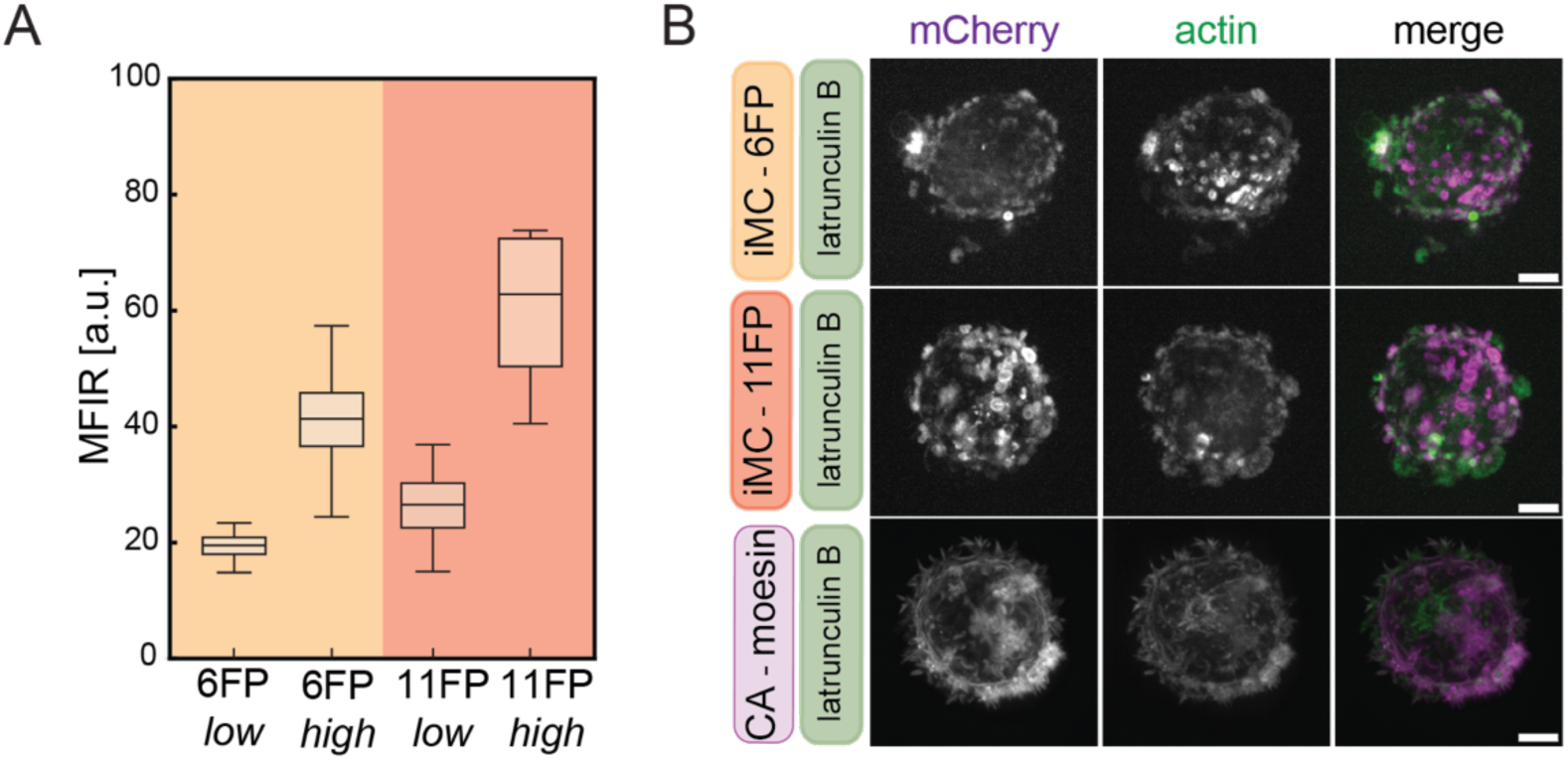
A) Boxplot measuring the mean fluorescence intensity ratio (MFIR) using flow cytometry of single cell clones with low and high expression levels of iMC-6FP and iMC-11FP. B) Representative images of the distribution of CA-Moe, iMC-6FP, and iMC-11FP in cells treated with 500 nM Latrunculin B. Scale bars = 5 μm. a.u. = arbitrary units. Solid line in box plot represents median.

**Figure S8.**
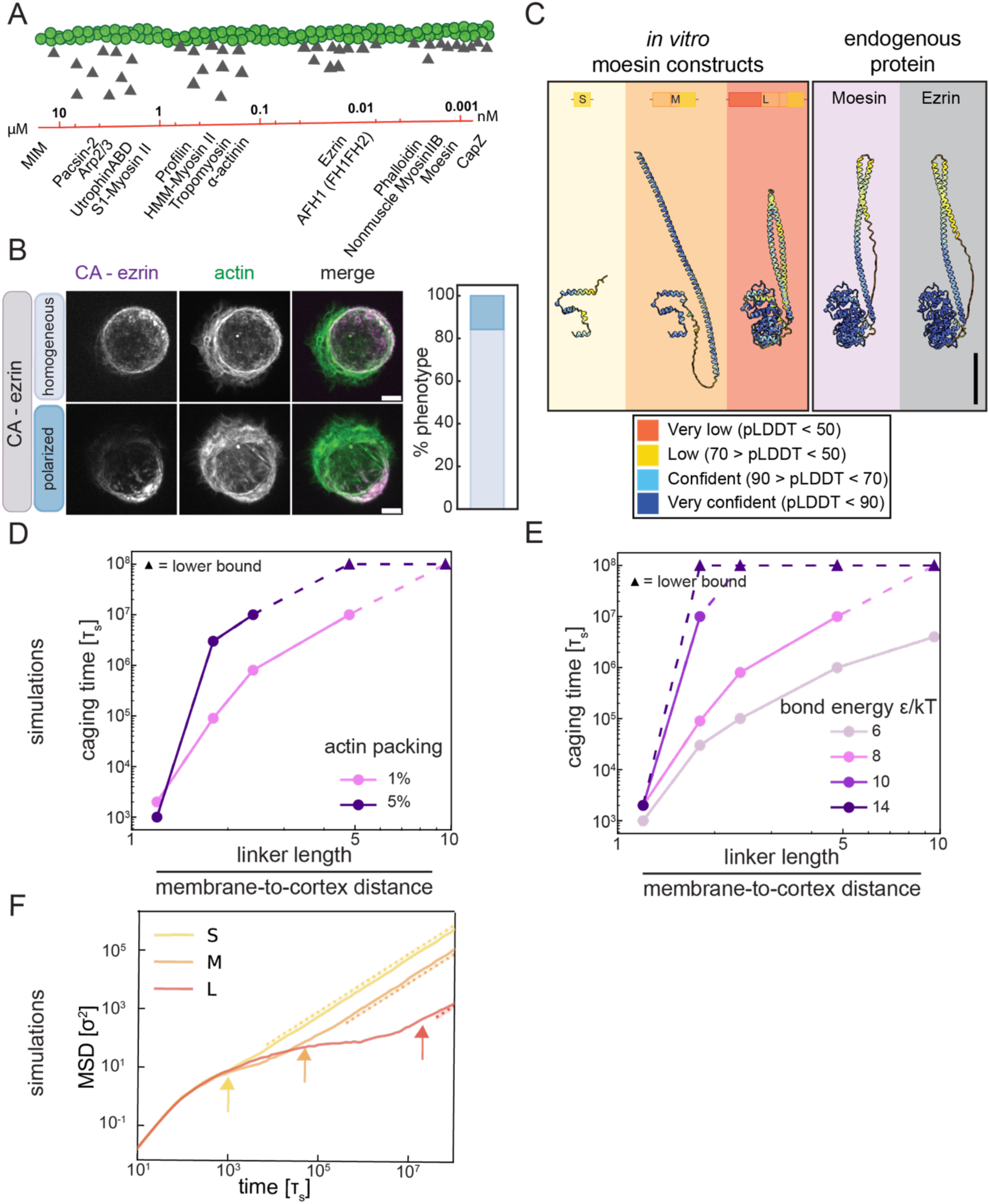
A) An affinity scale of F-actin binding proteins as reported in literature. Triangles on filament are a visual aid of the binding kinetics, see also Table S3. B) Images of NIH3T3 fibroblasts overexpressing the phosphomimetic version of ezrin (mouse), along with frequency of phenotypes (N = 3 experiments, 46 cells). C) AlphaFold predictions of phosphomimetic *in vitro* moesin constructs and endogenous non-phosphorylated proteins. Note that in predictions ezrin and moesin the actin binding and the plasma membrane binding domains are interacting and thus do not depict the fully extended protein. D-F) Caging time from simulations plotted as a function of actin packing (D) and bond energy (E). F) Actin mean square displacement from simulations. Arrows define the caging times. Straight dashed lines, slightly displaced for better readability, are a fit representing diffusive behaviour. Scale bars = 5 μm.

