## Supplementary tables 1-3 for "Caging of membrane-to-cortex attachment proteins can trigger cellular symmetry breaking"

| | Membr+M-Moe | | | Membr+MAC | | |
|---|---|---|---|---|---|---|
| | Recovered-100% | Recovered <100% | No Recovery | Recovered-100% | Recovered <100% | No Recovery |
| DOPC | 6 | 11 | 0 | 5 | 20 | 1 |
| Chol | 2 | 13 | 0 | 0 | 14 | 10 |
| DOPE | 3 | 10 | 0 | 1 | 12 | 8 |

Table S1: Frap recovery curves distribution for GFP-M-moesin on DOPC, Chol and DOPE membranes

| | Membr+S-Moe | | | Membr+MAC | | |
|---|---|---|---|---|---|---|
| | Recovered -100% | Recovered <100% | No Recovery | Recovered -100% | Recovered <100% | No Recovery |
| DOPC | 2 | 16 | 0 | 2 | 18 | 0 |
| Chol | 4 | 11 | 0 | 0 | 15 | 1 |
| | Membr+L-Moe | | | Membr+MAC | | |
| | Recovered -100% | Recovered <100% | No Recovery | Recovered -100% | Recovered <100% | No Recovery |
| DOPC | 2 | 18 | 0 | 0 | 14 | 8 |

Table S2: Frap recovery curves distribution for GFP-S and L-moesin on DOPC and Chol membranes

| Protein | Kd | Reference |
| --- | --- | --- |
| CapZ | $0.08 \pm 0.02\ nM$ | Schafer et al., *J Cell Biol* (1996). 135 (1): 169–179 |
| Moesin | $1.5\ nM$ (1.8 μM) | Nakamura et al., Mol Biol of the Cell (1999). 10: 2669–2685; Simons et al., Biochemical and Biophysical Research Communications (1998) 253(3):561-565 |
| Non-nuscle MyosinIIB (S1) | $3.2 \pm 0.02\ nM$ | Wang et al., *J Cell Biol* Chem (2003). 238(30): 27439–27448 |
| Phalloidin | $9 \pm 2\ nM$ | De La Cruz and Pollard. Biochemistry (1996). 35: 14054-14061 |
| AFH1 (FH1FH2) C-terminal | | Michelot et al., The Plant Cell (2005) 17: 2296–2313 |
| Ezrin | $50\ nM$ (5 μM) | Yao et al., *J Cell Biol* Chem (1996). 271(12): 7224–7229; Simons et al., Biochemical and Biophysical Research Communications (1998) 253(3):561-565 |
| α-actinin | 0.4 μM | Wachstock et al., (1995). BiophysJ 65:205-214 |
| Tropomyosin | 0.48 μM | Racca et al., J Biol Chem (2020). 295 (50): P17128-17137 |
| HMM- Myosin II | 0.58 μM | Miyata et al., J Biochem (1989) 105: 103-109 |
| Profilin | 1 μM | Courtemanche and Pollard. Biochemistry (2013). 52(37) |
| Pacsin2 | 2 μM | Salzar et al., Cell Mol Life Sci.(2017) 74(13): 2413–2438. |
| Arp2/3 | ~3-4 μM | Gournier et al., Molecular Cell (2001). 8 1041–1052 |
| S1-Myosin II | 11 μM | Racca et al., J Biol Chem (2020). 295 (50): P17128-17137 |
| MIM | 17 μM | Salzar et al., Cell Mol Life Sci.(2017) 74(13): 2413–2438. |

Table S3: An affinity scale of F-actin binding proteins as reported in literature
